# Post-transcriptional dysregulation precedes morphological transformation in chemically induced liver cancer

**DOI:** 10.64898/2026.09.30.755803

**Authors:** Osagie G. Izuogu, Michael Jackson

**Affiliations:** Dysplasia Diagnostics Limited, Cambridge, UK; Institute of Genetic Medicine, Newcastle University, UK

## Abstract

DNA lesions impede RNA polymerase II, yet the transcriptional consequences of that stress are less understood than the lesions themselves. We reanalysed total RNA and matched whole genomes from 438 liver lesions across four divergent mouse backgrounds, one of them the separate species *M. caroli*, 428 of them induced by a single dose of diethylnitrosamine (DEN) and 10 arising spontaneously, together with untreated liver and, critically, histologically normal liver from treated, tumour-bearing animals (paired normal liver).

Three post-transcriptional changes accompany transformation in every background. Canonical splicing efficiency falls by 3.6 to 4.8 percentage points, and in genes that acquire them, cancer-specific junctions account for a median of 20–25% of junctions and arise at weaker splice signals. Output shifts toward circular RNA: a five-method consensus pipeline identifies 56,418 recurrent backsplice junctions, 9,718 of them detected in all four backgrounds, with ∼2-fold enrichment in circRNA abundance and an expanded repertoire of distal configurations. Adenosine-to-inosine editing sites fall 2.2-to 3.8-fold, most steeply at intronic sites.

Exposure changes not only these outputs but their coordination. In untreated C57BL/6J liver, splicing efficiency, mean relative backsplice span and editing co-vary positively (pairwise ρ +0.78 to +0.87). That coordination is lost in exposed tissue, and paired normal liver shows the same loss as tumour. Pooling conditions obscures the effect because opposing estimates cancel.

In C57BL/6J, paired normal liver already shows tumour-like splicing, circRNA abundance and editing roughly eight months after one exposure, while backsplice architecture has begun to reorganise. These samples carry a median of 337 callable somatic single-nucleotide variants against 78,843 in tumours and show no elevation of *Mki67* or cell-cycle programmes. The post-transcriptional phenotype is therefore a durable property of damaged tissue established before morphological transformation.

## Introduction

Many cancers are initiated by unrepaired DNA damage that produces deleterious genetic aberrations. Mammalian cells encounter DNA lesions from endogenous and environmental agents that trigger conserved high-fidelity DNA repair processes to preserve genome integrity^1–4^. Lesions that persist through replication are converted into mutations, some of which contribute to transformation, and their accumulation generates the genetic heterogeneity of tumours^5^.

The effects of DNA damage on genome stability have been studied extensively, revealing the intricacies of damage sensing, cell-cycle control, repair and apoptosis^3,6–9^. Each iteration of transcription proceeds through recruitment of transcription factors, initiation and elongation, co-transcriptional RNA processing and termination^10^. Gene expression is context-dependent and tightly regulated to maintain a balance in the expression of all products from each locus. Several protein-coding genes with established oncogenic activities are frequently dysregulated in cancer, and there is accumulating evidence that non-coding RNA species have functional roles in gene regulatory circuits. Sam68, a splice factor, induces changes in Bcl-x isoform usage that promote resistance to apoptosis^11^; circular RNAs (circRNAs) act as competing endogenous RNAs that sequester RNA-binding proteins and miRNAs, with consequences for gene expression in cancer^12–16^, and exon-intron circRNAs can promote transcription of their host genes in the nucleus^17^.

DNA lesions alter transcriptome composition by several routes. Transcription-blocking lesions trigger transcriptional arrest to initiate transcription-coupled repair^18,19^, producing truncated transcripts^20^. Trans-lesion synthesis generates mutant transcripts with mis-incorporated nucleotides^21^, potentially affecting RNA–RNA and RNA–protein interactions. Direct chemical modification of RNA nucleobases impedes translation and reduces protein levels^22–24^. And to prevent transcription–replication collisions mediated by R-loops^25,26^, stalled RNA polymerase II (RNAPII) is dislodged and its largest subunit, RPB1, degraded, depleting the polymerase pool^27,28^.

These routes are usually considered one at a time, and each is framed as an acute, lesion-level event. But co-transcriptional RNA processing is kinetically coupled to transcription: splice-site recognition competes for a finite window while polymerase transits the gene, and the output of a locus depends on the relative rates of elongation and processing^15,29–32^. A tissue in which that coupling has been perturbed should show not one defect but a coordinated shift, canonical junctions spliced less efficiently, cryptic junctions used more often, and a greater share of nascent transcript diverted into circular forms. Whether such a shift occurs *in vivo*, how long it outlasts the initiating exposure, and whether it depends on the mutations that ultimately define a tumour, have not been established.

Circular RNA and RNA editing bring biology of their own to this picture. Spliceosome-mediated circRNAs arise from backsplice events joining downstream splice donors to upstream splice acceptors, forming covalently closed molecules ∼3-fold more stable than cognate linear RNAs owing to resistance to exoribonuclease activity^33,34^. Adenosine-to-inosine editing, catalysed by adenosine deaminase (ADAR) proteins, enhances transcriptome and proteome diversity^35–38^; ADARs also regulate R-loop formation^39^, influence circRNA biogenesis by destabilising double-stranded RNA^37,40^ and influence other co-transcriptional processes, including splice code modification^36,41,42^. Editing of repeat-derived duplexes in an intron can occur only while that intron is present in the nascent transcript, so intronic editing should also depend on the window available for co-transcriptional processing.

The diethylnitrosamine (DEN) model suits this question. DEN is a genotoxic carcinogen that requires metabolic activation in the liver, which is why the liver is its target^43^, and a single dose produces a well-characterised mutational signature^5,44^, so exposure is fixed in time and the damage it causes can be read from each lesion’s genome. Susceptibility to DEN and tumour latency differ markedly between mouse backgrounds^43,45^, so treating several genetically divergent backgrounds identically separates responses general to exposure from those peculiar to one genome.

The cohort analysed here was generated by the Liver Cancer Evolution Consortium and is described in full by Aitken et al.^45^, whose analyses address genome evolution, mutational processes and the genetic control of tumour trajectory. Here we use its RNA data to examine co-transcriptional processing and, critically, use histologically normal liver collected from treated animals (hereafter paired normal liver) to ask when the resulting molecular state becomes established relative to morphological transformation. We undertake an integrated analysis of mouse liver cancer genomes and transcriptomes to assess the cascading molecular events that accompany exposure to a carcinogen. Our results describe extensive changes to co-transcriptional splicing that are independent of mutations in cis-acting elements, increased circRNA formation with an expanded repertoire of distal backsplice configurations, and specific reductions in intronic RNA editing associated with *Adarb1* transcript usage, and a loss of the coordination that links splicing efficiency, backsplice span and editing in untreated liver. We find these changes in paired normal liver, months after a single exposure and in the absence of both the mutation burden and the proliferation that characterise the lesions themselves.

## Results

### Transcriptome diversity is enhanced in transformed tissue

We analysed the transcriptomes of a cohort in which early liver cancer evolution was re-run across four divergent mouse backgrounds, *Mus musculus domesticus* (C57BL/6J and C3H/HeOuJ), *M. m. castaneus* (CAST/EiJ) and *M. caroli* (CAROLI/EiJ), following a single intraperitoneal dose of DEN at fifteen days of age. That cohort, its treatment protocol, histopathology, genome sequencing and driver landscape are described in full elsewhere^5,44,45^; what follows is a post-transcriptional analysis of the same material.

In brief, tumours were harvested at background-specific latencies and comprise dysplastic nodules and hepatocellular carcinomas spanning different stages of transformation, a subset of which arose spontaneously in untreated animals. Matched whole genomes and ribosome-depleted total RNA were prepared from the same tissue. Alongside the tumours we analyse untreated liver at postnatal day 15 (P15, the age of exposure) and untreated adult liver age-matched to tumour isolation^45^ and, critically, twelve paired normal samples, histologically normal liver harvested from DEN-treated, tumour-bearing C57BL/6J animals, which are the informative group for the timing argument developed below (Fig. 1a). We use lesion for an individual dysplastic nodule or hepatocellular carcinoma, and tumour for lesions as a group in comparisons with normal liver.

**Figure 1.**
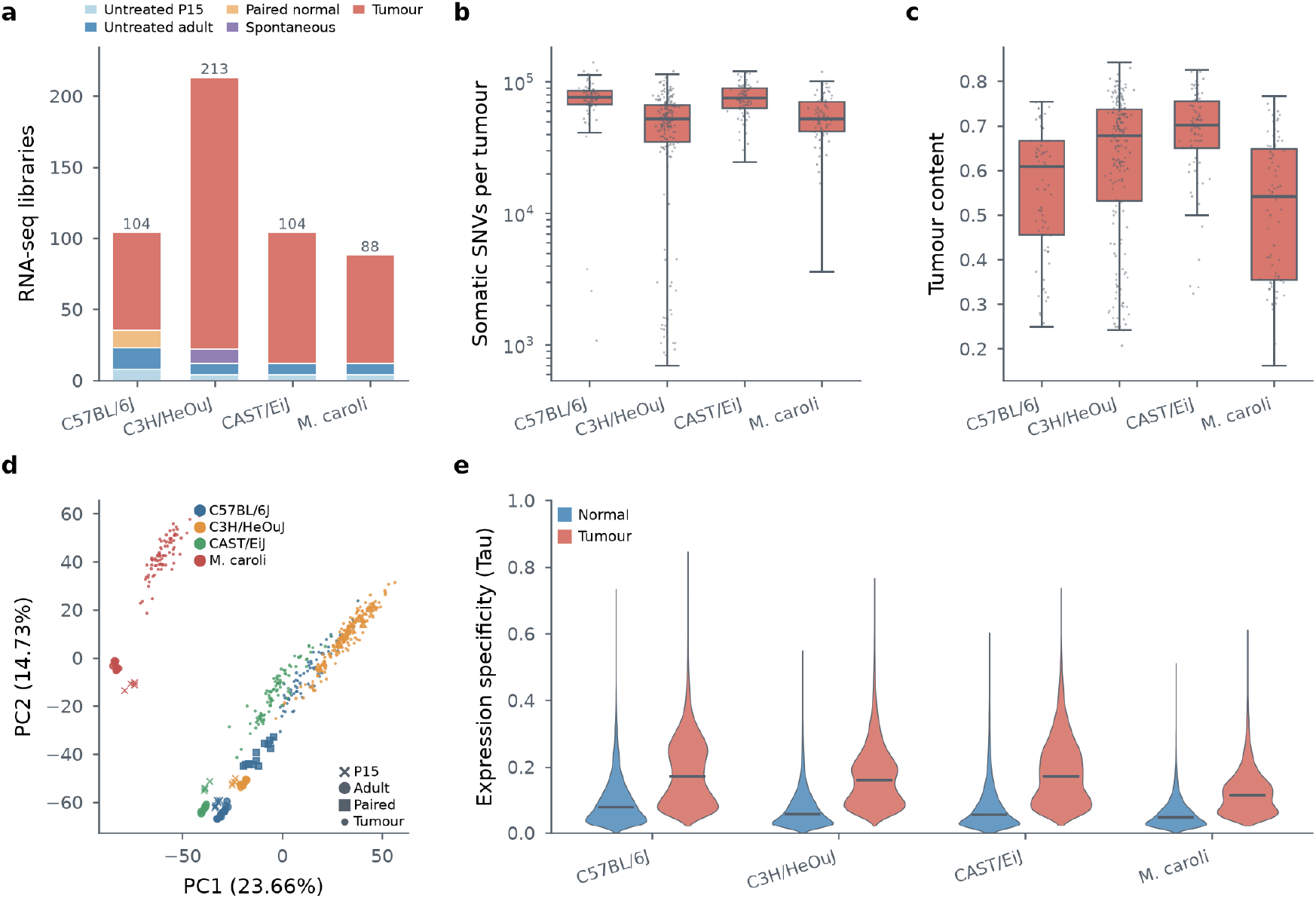
The cohort, and how transcriptomes differ across it. **a**, RNA-seq libraries by background and sample class. The asymmetry matters for everything that follows: histologically normal liver from DEN-treated animals exists for C57BL/6J and essentially nowhere else, and the spontaneous lesions are C3H/HeOuJ only. **b**, Somatic single-nucleotide variants per lesion, by background, log scale. Boxes are quartiles with the median; points are lesions. **c**, Tumour content per lesion, as the genomic cellularity estimate (ploidy x modal variant allele fraction; see Methods). **d**, First two principal components of expression across all four backgrounds, on the 1,000 most variable genes of the variance-stabilised matrix, coloured by sample class. **e**, Expression specificity (Tau) in normal liver against tumours, by background, from the cohort’s own per-gene table. Tau is the Yanai index, 0 where a gene is expressed evenly across samples and approaching 1 where it is confined to one. Medians rise from 0.048-0.078 in normal liver to 0.114-0.171 in tumours, and 91-97% of individual genes increase: expression becomes lesion-restricted rather than uniform. Tau’s absolute value depends on the expression transform it is computed from, so the published values are used rather than a recomputation.

Whole-genome sequencing of the same lesions reports between 3,630 and 140,579 mutations per lesion, with the highest genome-wide burdens in C57BL/6J (median 78,843; Fig. 1b)^5^. Paired normal samples carry between 66 and 3,956 mutations and no recurrent driver mutations, the early mutational consequences of exposure in histologically normal tissue. Tumour cell content, estimated from ploidy and the modal variant allele fraction, spans 0.16 to 0.84 across the 438 lesions (median 0.64; Fig. 1c), and is carried as a covariate where biologically and statistically appropriate.

To examine the effects of DNA damage on transcriptome composition we profiled Illumina ribosome-depleted total RNA-seq datasets, with on average over 26 million uniquely mapped reads per sample, ∼75% of the reads sequenced (reads mapping to up to ten loci were additionally retained; Methods). Read-distribution analysis showed consistent differences in the proportion of reads originating from well characterised gene biotypes (Supplementary Fig. 1a–d). Globally, the proportion of reads mapping to protein-coding genes, long non-coding RNAs and host genes of small non-coding RNAs is reduced in tumours relative to untreated liver. Conversely, reads originating from processed pseudogenes increase, consistent with reported activation of pseudogene transcription in cancer^46^. Low-dimensional representations of gene expression using principal components analysis separate samples by condition, background and stage of transformation (Fig. 1d).

We investigated changes in protein-coding gene expression specificity and consistency using the Tau method^47^. We observed reproducible shifts in expression variability, from broadly expressed levels in normal samples to oscillating, nodule-specific levels for the same genes in tumours (Fig. 1e). Comparing each gene with itself, Tau rises in 91.4% of genes in C57BL/6J and in 96.5%, 95.9% and 94.0% in C3H/HeOuJ, CAST/EiJ and *M. caroli*, and the median roughly doubles to trebles in every background (all genes; C57BL/6J 0.078 to 0.171, C3H/HeOuJ 0.057 to 0.159, CAST/EiJ 0.056 to 0.171, *M. caroli* 0.048 to 0.114; paired Wilcoxon *P* < 1 × 10^−300^ throughout, 16,797–19,450 genes per background). The shift is a property of the transcriptome rather than of a subset of genes large enough to move a distribution: virtually every gene expressed in this tissue is read out in a lesion-restricted way that it is not in normal liver.

The scale of that shift is easiest to read against the non-coding transcriptome, where restricted expression is the norm rather than the exception. In C57BL/6J, the median protein-coding gene in tumours reaches a specificity of 0.149 against 0.100 for non-coding genes in normal liver, and 69.7% of protein-coding genes exceed that non-coding baseline; in *M. caroli* the same comparison gives 0.108 against 0.049 and 91.2%. Protein-coding transcription in tumours is therefore distributed across samples more like non-coding transcription in normal liver than like its own behaviour there. Differential expression analysis, untreated adult liver against tumour within each background, identified between 8,757 (CAST/EiJ) and 10,507 (C57BL/6J) significantly altered protein-coding genes (C3H/HeOuJ 9,379, *M. caroli* 9,518; adjusted *P* < 0.05), of which 3,627–4,268 change by more than two-fold. Tested against the genes each analysis could detect, genes higher in tumours are enriched in every background for translation, biosynthetic processes and protein transport, and for mitotic cell-cycle processes, while genes lower in tumours show little coherent enrichment (Supplementary Data 1).

At the gene level, nearly every core RNA polymerase II subunit is dysregulated in tumours. In C57BL/6J tumours, *Polr2a*, encoding the largest RNAPII subunit (RPB1), is ∼4-fold lower than in untreated liver while ten of the remaining eleven core subunits are higher (Supplementary Fig. 1e,f). *Polr2a* expression is known to be altered in response to DNA damage^28^; here it remains repressed about eight months after a single DEN dose, in contrast to reports of post-repair recovery^26,48^. This is the first indication in these data that the transcriptional consequences of a single exposure are not transient.

Collectively, despite differences in genetic background and tumour latency, we find recurrent alterations to transcriptome composition in all four backgrounds.

### Canonical splicing efficiency is reduced in tumours

Splice isoform variation has been reported in cancer initiation, progression and resistance to therapy^49–51^. We assessed splicing efficiency in two ways. First, for each gene we derived a Splicing Efficiency Index (SEI) by dividing the number of spliced reads at the donor and acceptor sites of Ensembl-annotated^52^ splice junctions by the total number of spliced and unspliced reads overlapping those sites in each sample (Methods; Supplementary Data 2). Unspliced reads at splice junctions typically originate from nascent, truncated, incompletely processed or alternatively spliced transcripts comprising retained introns or bleeding exons processed at cryptic splice sites^53^. In untreated liver the per-sample mean SEI across genes exceeds 86%, providing an indication of nascent transcript levels in the sequenced libraries.

We characterised the sampling distribution of mean SEI for each condition (Fig. 2a). The distributions are bimodal in untreated liver, resolving the two ages sampled: mean SEI is 86.7% for P15 and 88.1% for adult C57BL/6J liver, adult liver being the more efficient. Adult liver exceeds P15 in the other three backgrounds too, by 1.11, 1.09 and 0.84 percentage points in C3H/HeOuJ, CAST/EiJ and *M. caroli*, consistent with age-related differences in co-transcriptional splicing.

**Figure 2.**
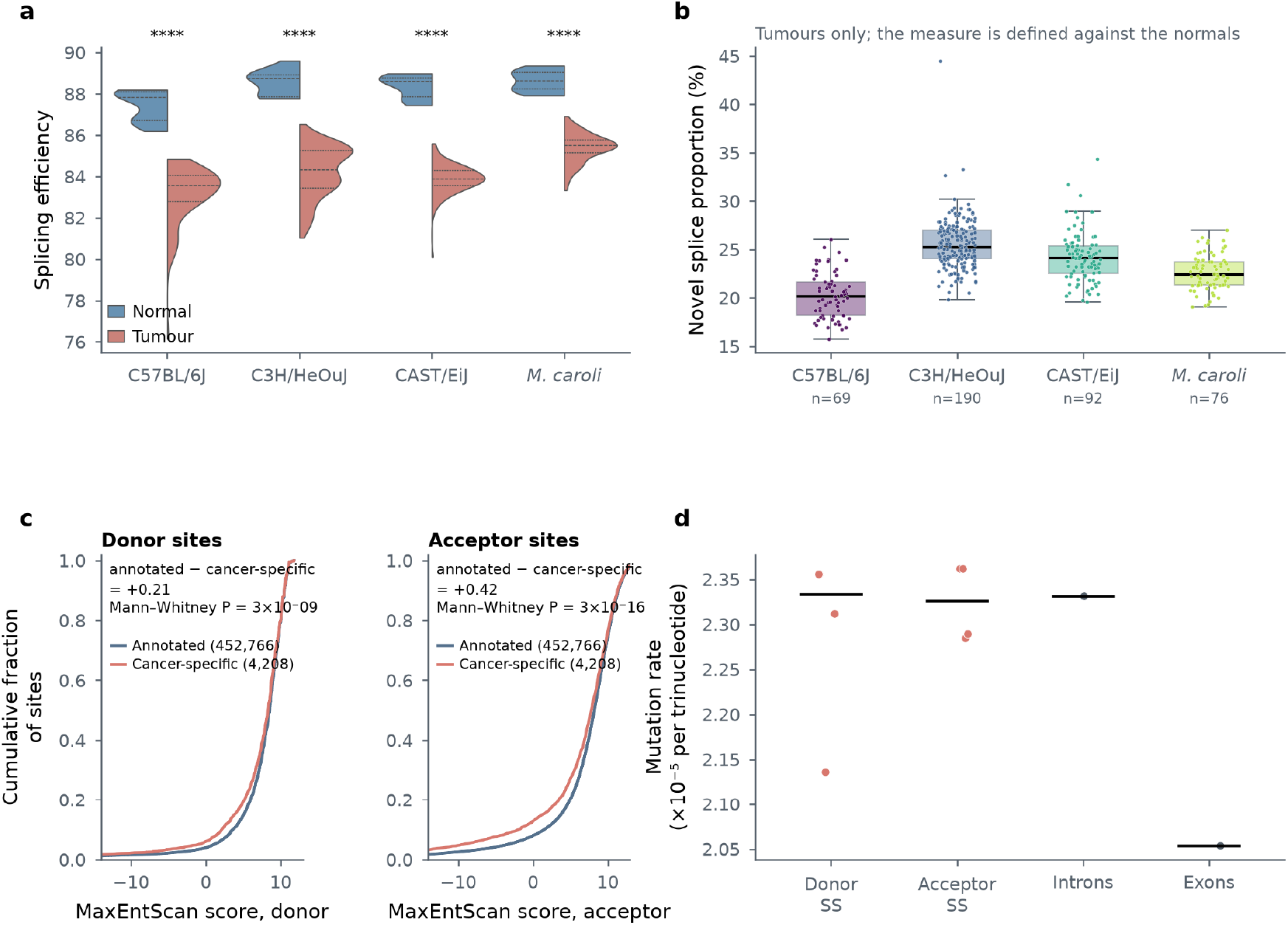
Canonical splicing efficiency falls and cryptic splice-site use rises. **a**, Splicing efficiency by background, untreated liver against tumours, as split violins with quartile lines. The untreated distribution is bimodal in each background; the two modes are P15 and adult liver, so the shape carries the age structure and the ages are not broken out here (the text gives the two modes). Mann–Whitney per background. **b**, Novel splice proportion in tumours. The measure counts junctions absent from the normals, so it is undefined outside tumours and is shown by background rather than by condition. **c**, MaxEntScan splice-signal strength for annotated against cancer-specific sites, as cumulative distributions. Cumulative rather than histogram form because the two sets differ a hundred-fold in size and the shift sits in the lower tail. Donor Δmedian +0.21, acceptor +0.415. **d**, Somatic mutation rate by feature class, trinucleotide-normalised on the scale used for this cohort^5^. Splice sites sit at the intronic rate; exons are the one depleted class, which follows from transcription-coupled repair rather than from any property of splice sites. Points are the subsets recorded in the source tables; bars are their medians.

The P15-to-adult difference is ∼1.4 percentage points in C57BL/6J, smaller than any untreated-to-tumour difference (below), and that gap does not scale with the age at which animals were sacrificed: *M. caroli*, sampled latest (56–80 weeks after exposure, median 80, against 22–39 weeks, median 25, for C3H/HeOuJ) and known to evolve cancer most slowly^45^, shows the smallest gap, the reverse of the ordering chronological age would predict. Tumours have significantly lower SEI than untreated adult liver in every background (C57BL/6J −4.5, C3H/HeOuJ −4.6, CAST/EiJ −4.8, *M. caroli* −3.6 percentage points; all *P* ≤ 1.3 × 10^−6^, and every interval excludes zero when animals rather than lesions are resampled; Fig. 2a), indicating alterations to the splicing programme. Per gene, the standard deviation of SEI across tumours has a median of 4.3–5.3 percentage points, and it is associated with expression specificity: a linear model of Tau on SEI variation explains 31–38% of its variance across genes (R² per background); because the two measurements share reads, this should be read as an upper bound.

### Cryptic splice-site use rises as canonical use falls

To estimate cancer-specific splicing deregulation directly, we calculated the Novel Splice Proportion (NSP) as the proportion of cancer-specific splice junctions in each gene. In genes carrying at least one cancer-specific junction, such junctions account for a median of 24.0% of the gene’s junctions across tumours, from 20.1% in C57BL/6J to 25.2% in C3H/HeOuJ (Fig. 2b); averaged over all genes, including those without one, the figure is 1.7–3.1%. Clone type, which indicates tumour cell origin, is associated with NSP in one background. Every C57BL/6J lesion is clonal, so the comparison cannot be made there, and there is no association in C3H/HeOuJ (Kruskal-Wallis *P* = 0.35) or CAST/EiJ (*P* = 0.54). In *M. caroli*, whole-genome-duplicated lesions have lower NSP than clonal lesions (medians 21.5% and 23.4%; *P* = 4 × 10^−4^ across the three clone types), a difference that holds when animals rather than lesions are resampled (−1.9 percentage points, 95% CI −2.5 to −0.9).

As cryptic splice sites are typically activated during cellular stress and are otherwise blocked by RNA secondary structure or proximity to well positioned splice sites^54,55^, we assessed whether splice signal strength contributes to the observed deviations. Genomic sequences of donor and acceptor regions for both annotated and cancer-specific sites were examined using MaxEntScan^56^. In all four backgrounds, cancer-specific donor and acceptor sites scored lower than annotated sites. In C57BL/6J the medians differ by 0.21 and 0.415 MaxEntScan units (8.55 against 8.34 at donors and 8.21 against 7.795 at acceptors; *P* = 3 × 10^−9^ and 3 × 10^−16^, Mann–Whitney, 452,766 annotated against 4,208 cancer-specific sites; Fig. 2c), and by 0.05–0.25 units at donors and 0.19–0.53 at acceptors in the other three, significantly in CAST/EiJ and *M. caroli* and not in C3H/HeOuJ, where 1,149 cancer-specific donor sites were recovered against 6,315 and 4,480 (Supplementary Fig. 2), consistent with increased use of weaker sites under altered splice-site competition rather than with acquisition of strong new sites.

To identify functional categories of genes with tumour-specific splice patterns, we analysed samples using rMATS^57^, screening for statistically significant alternatively spliced events (ASEs). We identified 2,482 (C57BL/6J), 5,154 (C3H/HeOuJ), 4,518 (CAST/EiJ) and 4,334 (*M. caroli*) significant events at FDR < 0.05 with greater than 10% inclusion-level difference relative to normal samples (Supplementary Data 3). In all four backgrounds the majority are mutually exclusive exon events, 1,927, 4,649, 3,799 and 3,534 respectively, against 299, 214, 315 and 412 skipped exons, 60, 94, 140 and 105 alternative 3′ splice sites, 44, 48, 72 and 77 alternative 5′ splice sites, and 152, 149, 192 and 206 retained introns. This identified a population of genes undergoing extensive alternative splicing in tumours in all four backgrounds. Testing those genes against the set of genes rMATS could test in the same background, rather than against the genome, they are enriched for the metabolic and secretory programmes that define hepatocyte identity: xenobiotic metabolism, fatty acid metabolism, bile acid metabolism, peroxisomal function, adipogenesis and coagulation reach significance in all four backgrounds, and oxidative phosphorylation, protein secretion and complement in three (Supplementary Data 1). Two growth-signalling programmes are also enriched, mTORC1 in all four backgrounds and PI3K/AKT/mTOR in three. Recurrent alternative splicing therefore affects predominantly liver metabolic and secretory programmes, alongside reproducible enrichment of mTOR-related growth signalling.

### Splice-region mutations do not explain the change

Genetic mutations proximal to splice sites may enhance or repress utilisation. Canonical splicing efficiency shows a weak negative association with DEN signature exposure in *M. caroli* (ρ = −0.306, *P* = 0.007, n = 76), in the direction predicted if damage-associated exposure depressed splicing, but no comparable association in the other three backgrounds, and it does not survive correction for the number of background-by-measure combinations tested. The second DEN signature component runs in the opposite direction in the same background (ρ = +0.240), which is difficult to reconcile with a single mechanism. We then assessed mutation burdens in 1,000-bp flanking regions of splice sites observed in both normal and tumour samples of a background (common) and those observed only in tumours (tumour-only), and compared these with genome-wide mutation rates. Although there are observable differences between strains, rates at splice regions are at or below the intronic rate wherever introns were measured (Fig. 2d; Supplementary Fig. 2c,f,i). Genome-wide rates here are trinucleotide-normalised, mutations in a given trinucleotide context divided by that context’s count in the genomic space assessed, following the framework established for this cohort^5^ and used subsequently to resolve mutagenicity by strand^58^.

Comparing feature classes directly on that scale, splice sites are not preferentially mutated. In C57BL/6J, donor sites carry rates of 2.14–2.36 × 10^−5^ and acceptor sites 2.28–2.36 × 10^−5^, against 2.33 × 10^−5^ across introns and 2.05 × 10^−5^ across exons. Splice sites therefore sit at the intronic rate rather than above it, and the one class that is measurably depleted is exons. The ordering holds elsewhere (Supplementary Fig. 2c,f,i): splice sites sit at or below the intronic rate wherever introns were measured, 2.48 and 2.53 against 2.78 × 10^−5^ in CAST/EiJ and 1.98 and 1.96 against 2.25 × 10^−5^ in *M. caroli*, and exons are the lowest class in C3H/HeOuJ and CAST/EiJ. Only in *M. caroli* are the three indistinguishable, exons at 1.97 × 10^−5^ against 1.96–1.98 at splice sites. That ordering is expected rather than surprising: mutation rate in this cohort falls with transcription, and most steeply in the most highly expressed genes^58^, so the depletion tracks transcriptional activity rather than any property of splice sites themselves. Mutations within 10 bp of splice sites represent on average <10% of mutations in each tumour.

To identify variation in gene expression attributable to splice-region mutations, we performed regression analysis relating the observation of any splice-region mutation in each gene to expression estimates across samples, controlling for tumour cell content and driver gene. This is analogous to splice-eQTL analyses^59^. After correcting for multiple testing, expression levels of 5, 6, 2 and 17 genes were significantly associated with splice-region mutations in C57BL/6J, C3H/HeOuJ, CAST/EiJ and *M. caroli* respectively, representing under 0.6% of genes tested in every background (Supplementary Fig. 3). Eleven of the seventeen genes in *M. caroli* lie within two adjacent stretches of its annotation, so even that background’s excess reflects a single divergent region rather than independent associations. Splice-region indels are too rare to test at comparable breadth, with only 24 to 234 genes reaching the minimum in any background, and they yield 1 to 32 genes each. Intratumour heterogeneity, measured as mutant-allele tumour heterogeneity (MATH) from whole-genome variant allele fractions, likewise shows no association with any post-transcriptional measure in any background once tumour cell content is accounted for (|ρ| ≤ 0.23, all *P* > 0.05).

The transcriptome-wide splicing phenotype described above cannot be explained by mutation of the cis-acting elements that specify splicing. The phenotype is therefore more consistent with trans-acting regulation than with recurrent mutation of the splice-site substrate; altered processing kinetics and trans-factor availability are candidate mechanisms. As concentrations of RNA-binding proteins, other trans-acting elements and mutations elsewhere in these genes may contribute to expression variability, the small number of associations cannot be attributed uniquely to splice-region mutations and may include contributions from other local or trans-acting factors.

### Output shifts from linear to circular RNA

Having identified a reduction in canonical splicing efficiency, we sought to characterise circRNA populations. Computational identification of backsplice junctions from RNA-seq can be confounded by several sources of artefact^60^. We developed a pipeline, ecircscreen, combining outputs of five published methods^12,60–63^ distributed across three aligners, and requiring each prediction to be generated by at least three of the five. Since no aligner carries more than two of the callers, this threshold guarantees corroboration across independent alignment strategies and so substantially reduces the aligner-specific biases that dominate differences between individual callers, before applying filters targeting known sources of false positive backsplice predictions. After removing predictions observed in single samples, predictions overlapping segmental duplications, and predictions supported only by suboptimally mapped reads, 56,418 distinct recurrent backsplice junctions were identified across all samples and placed on a common reference (Supplementary Data 4). Of these, 41,298 (73.2%) were detected in two or more backgrounds and 9,718 (17.2%) in all four, including *M. caroli*; 96% of junctions assignable to an annotated structure are exonic, rising to 99.3% among those detected in all four. An earlier implementation of ecircscreen was included as one of sixteen callers in an independent, developer-run community benchmark of circRNA detection^64^, where its conservative consensus and repeat filters place it toward the specificity end of the sensitivity-specificity trade-off, the appropriate operating point for the cross-background comparisons made here, in which a false-positive class shared across backgrounds would be indistinguishable from conservation. Matching on exact agreement at both breakpoints, approximately 71% are catalogued in circAtlas^65^ or circBase^66^, providing orthogonal support for circularity, and ∼55% match circRNAs previously reported from human tissues, so most are not specific to mouse. Both figures rise with cross-background recurrence, reaching 95% and 76% among junctions detected in all four backgrounds.

Grouping circRNAs by genomic region of origin across all per-background calls, including those not assignable to an annotated structure, >75% originate from exonic regions in every background (Fig. 3a); fewer than ∼20% originate from intronic or intergenic regions, consistent with previous reports^12,67^. More than 87% of backsplice predictions originate from protein-coding genes. Across all backgrounds the union of tumour libraries contains a broader repertoire of backsplice events than normal liver, and for virtually every circRNA host gene additional events are observed in tumours. Event discovery depends on the number and depth of libraries contributing to a group, so this is an expanded observed repertoire rather than a per-sample estimate of backsplice efficiency. What does change is which genes backsplice, and that is reproducible across backgrounds. Of the 7,686 genes producing circRNAs in tumours but not in normal liver of at least one background, 58.6% (4,504) do so in two or more backgrounds, 2,389 in three or more and 798 in all four (Fig. 3b); these reproducible host genes are significantly enriched in mitotic cell-cycle regulation, intracellular protein transport, RNA processing and cellular response to DNA damage (Supplementary Data 1).

**Figure 3.**
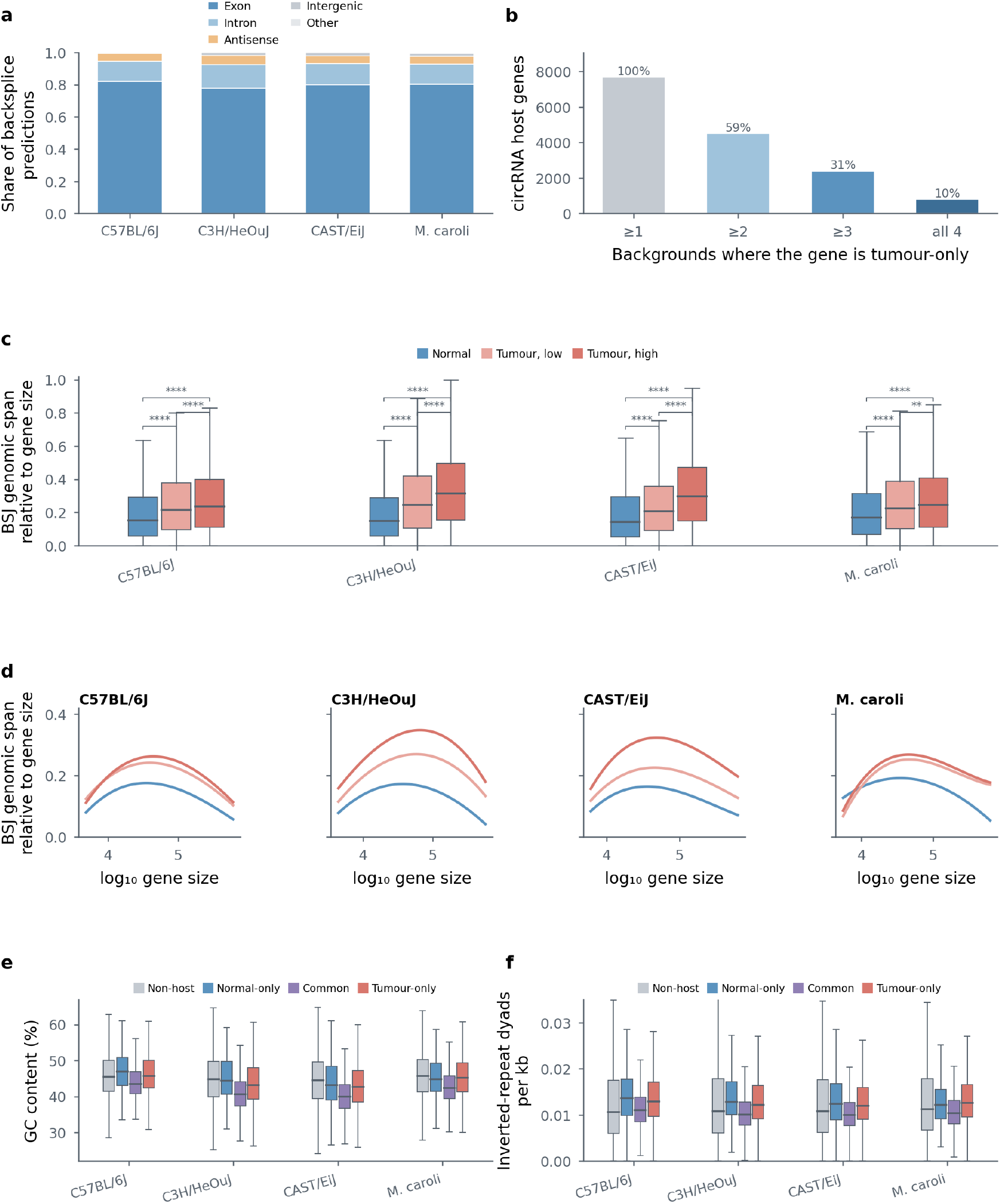
Output shifts from linear to circular RNA. **a**, circRNA biotype composition per background. Exonic circles comprise >75% of predictions in all four backgrounds, consistent with the expected biotype composition of circRNA catalogues. Recurrence across backgrounds, assessed on the projected common reference rather than on the per-background calls shown here, is not simply a function of depth: the proportion of junctions assignable to an annotated structure rises from 16.2% in one background to 96.3% in all four, and, among the pre-consensus candidate junctions, those called by all five callers are thirty-fold more likely to be conserved across all four backgrounds than those called by two. **b**, Reproducibility across backgrounds of genes producing circRNAs in tumours but not in normal liver; percentages are of the 7,686 genes tumour-only in at least one background. **c**, Condition-level maximal backsplice span relative to host gene size, for normal liver and for tumours split at cellularity 0.5, in all four backgrounds. Per gene the statistic is the maximum length-normalised span observed across the samples of that group, restricted to genes seen in more than one sample; paired normals are excluded and untreated P15 liver is the reference. Brackets are Mann–Whitney; ** *P* < 0.01, **** *P* < 0.0001. **d**, The same against host gene size, one panel per background. Bin medians over 33 bins of log₁₀ gene size, weighted by bin occupancy and smoothed with a cubic spline. The three conditions stay separated across the whole range, so the increase is not a consequence of tumours drawing circles from longer genes. **e**, GC content of circRNA host genes, partitioned by the conditions in which they circularise, against genes producing no circRNA. The GC-poor bias belongs to the constitutive set (medians 39.9–43.5%) rather than to host genes as a class; genes circularising only in tumours sit nearer the genome average (42.6–45.8% against 44.4–45.7% in non-host genes). Polymerase transits GC-poor sequence faster, so the constitutive set is the composition a kinetic account predicts and the tumour-specific set is not. **f**, Density of inverted-repeat dyads, the structural requirement for backsplicing, under the same partition and in the same direction. Mann–Whitney against the non-host reference.

The largest backsplice span observed per gene, relative to gene size, increases from normal liver through tumours of low and then high tumour content in all four backgrounds (C57BL/6J median 0.152, 0.216 and 0.235; C3H/HeOuJ 0.148, 0.245, 0.315; CAST/EiJ 0.143, 0.205, 0.297; *M. caroli* 0.168, 0.225, 0.245; all *P* < 2.2 × 10^−16^, Mann–Whitney; Fig. 3c; Supplementary Data 5). Because this statistic takes the maximum observed across all samples in a condition group, it describes the repertoire of backsplice configurations accessible to that condition rather than the typical span of an individual library: per-sample mean relative span shows no detectable adult-to-tumour shift, whereas cross-gene span dispersion does (see *The post-transcriptional state precedes transformation*; Supplementary Fig. 4). The separation holds across the whole range of host gene size in all four backgrounds (Fig. 3d) while contrasting with significant correlations with nucleotide composition; circRNA-producing genes have lower GC content than non-circRNA genes in every background, and partitioning host genes by the conditions in which they circularise locates that bias rather than spreading it over host genes as a class (Fig. 3e). It belongs to the genes that circularise in both conditions, the GC-poorest class in all four backgrounds (medians 39.9–43.5%, against 44.4–45.7% in genes producing no circRNA). Genes that circularise only in tumours sit between the two (42.6–45.8%), nearer the genome average than the constitutive host set. Inverted-repeat dyad density follows the same partition among host genes, lowest in the constitutive set and higher in both condition-specific sets (Fig. 3f).

Tumour backsplicing therefore recruits genes rather than intensifying an existing bias. The genes that circularise in normal liver are the ones sequence composition would predict: GC-poor, and transited faster by polymerase. The genes that begin circularising in tumours are not, and a mechanism acting through the substrate should have picked them out first. This is the same form of argument as the splice-region mutation result above, applied to a different process, and it points the same way: the change in exposed tissue argues against substrate sequence composition alone, and is consistent with a trans-acting change in RNA processing, including the kinetic account considered in the Discussion.

### circRNA enrichment is established before local morphological transformation

To assess circRNA expression patterns we performed hierarchical clustering using Spearman correlation coefficients of circular-to-linear junction count ratios, which grouped untreated liver samples by age and all samples by condition (Supplementary Fig. 5a). Within these groups, samples did not separate by tumour content, latency, driver gene or stage.

Comparing total backsplice junction (BSJ) expression per gene with cognate canonical splice junction (CSJ) expression, we observed the previously reported strong correlations that indicate shared mechanisms and origins^68^. However, when comparisons from both conditions were overlaid (Fig. 4a–c), expression patterns diverge: increased circRNA abundance corresponds to decreased canonical splicing expression for most genes in tumours. Comparing the log₂ change in backsplice against canonical junction expression between conditions (Fig. 4d–f; Methods) places the tumour-against-adult distribution at a backsplice log₂ change of ∼0.9 with canonical change centred on zero, a transcriptome-wide ∼2-fold enrichment in circRNA abundance in transformed tissue.

**Figure 4.**
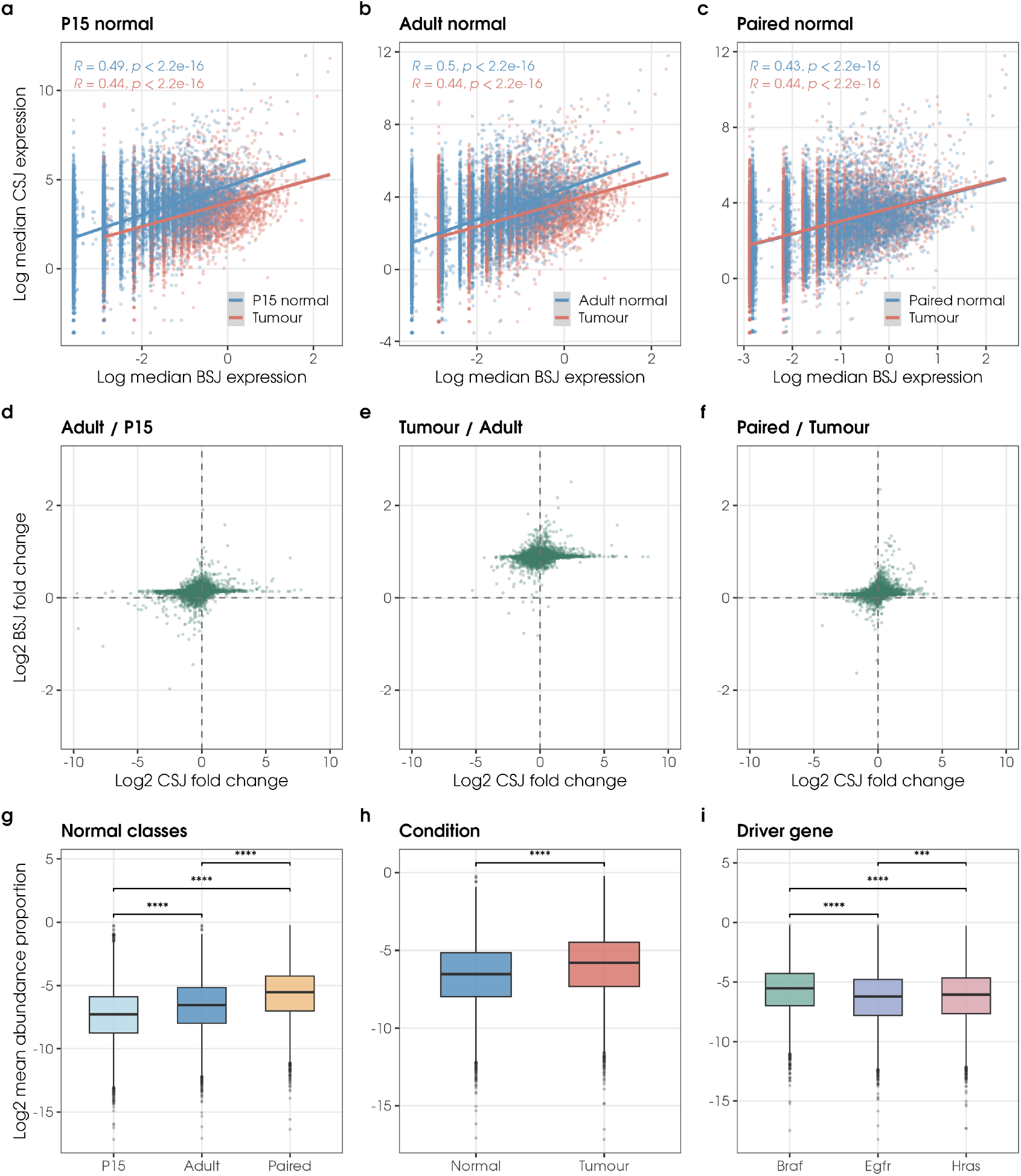
circRNA enrichment is established before local morphological transformation (C57BL/6J). **a**–**c**, Median canonical against median backsplice junction expression per gene, in junctions per million of total splice-junction reads, for each class of normal liver overlaid on tumour: untreated P15 (**a**), untreated adult (**b**) and paired normal liver (c). Lines are linear fits with 95% confidence bands; Pearson correlation inset. The two fits separate for both untreated classes and coincide for paired normal liver, which is the comparison the section rests on. **d**–**f**, Log₂ change in median backsplice against median canonical junction expression between adjacent classes. The tumour-against-adult distribution sits at a backsplice log₂ change of ∼0.9 with canonical change centred on zero, the transcriptome-wide ∼2-fold circRNA enrichment; adult-against-P15 and paired-normal-against-tumour sit at zero. **g**–**i**, circRNA abundance proportion, the ratio of backsplice to total spliced reads per gene, by class of normal liver (**g**), by condition (**h**) and by driver gene among tumours (**i**). Mann–Whitney; *** *P* < 0.001, **** *P* < 0.0001.

Strikingly, while these differences are apparent when P15 and adult normal samples are compared with tumours, there are no obvious differences in circRNA abundance profiles between tumours and paired normal samples. The same holds for editing (below). This is the central observation of this study and we treat it separately below (The post-transcriptional state precedes transformation). CircRNA abundance is marginally higher in adult than P15 normal samples, presumably reflecting age-related accumulation of circRNAs^68–72^. As a final confirmation of enrichment we compared circRNA abundance proportions (AP), the ratio of total BSJ read counts in each gene to total spliced reads from the same gene (Fig. 4g–i). AP distributions differ significantly between conditions (*P* < 2.2 × 10^−16^, Mann–Whitney) and are confirmed in cross-background comparisons. Grouped by driver gene, enrichment is significantly higher in samples with consequential *Braf* mutations, which transform earlier^5^; this was not consistent across strains, suggesting strain-specific effects or sampling bias. Because enrichment is already present in paired normal liver, which lacks recurrent drivers, driver mutations are not required for the early circRNA enrichment.

In highly proliferating cancer cells, steady-state circRNA levels are reported to be reduced, presumably diluted by increased cell division^73–75^. Our data are incompatible with dilution as an explanation, in both directions. Tumours proliferate, *Mki67* expression is elevated relative to both normal and paired normal liver (*P* = 4.1 × 10^−5^ and 7.3 × 10^−7^), *and* carry more circRNA, which is the opposite of the sign dilution predicts. Conversely, paired normal samples have the *lowest Mki67* of the three groups, indistinguishable from untreated normal liver (*P* = 0.19), yet carry near-tumour circRNA abundance profiles. Proliferation is therefore neither necessary for circRNA enrichment nor able to account for its direction. Unlike previous studies of established cancers, these data show circRNA enrichment already present in exposed tissue, before any reduction that may accompany later clonal expansion.

### A-to-I RNA editing sites are depleted in exposed tissue

We used SPRINT^76^ to profile editing events. After filters excluding predictions overlapping segmental duplications, homopolymer repeats and genome variants, we identified between 155,036 (CAST/EiJ) and 239,651 (C3H/HeOuJ) distinct editing events, the majority (∼87% in C57BL/6J) originating from interspersed and tandem repeats (Supplementary Data 6). Between 2.2- and 3.8-fold more editing sites (RES) were identified in untreated adult liver than in tumours (Fig. 5a): median counts fall from 9,898 to 4,048 in C57BL/6J (2.5-fold), 7,180 to 2,717 in C3H/HeOuJ (2.6-fold), 9,186 to 2,415 in CAST/EiJ (3.8-fold) and 7,296 to 3,386 in *M. caroli* (2.2-fold), relative to untreated adult liver.

**Figure 5.**
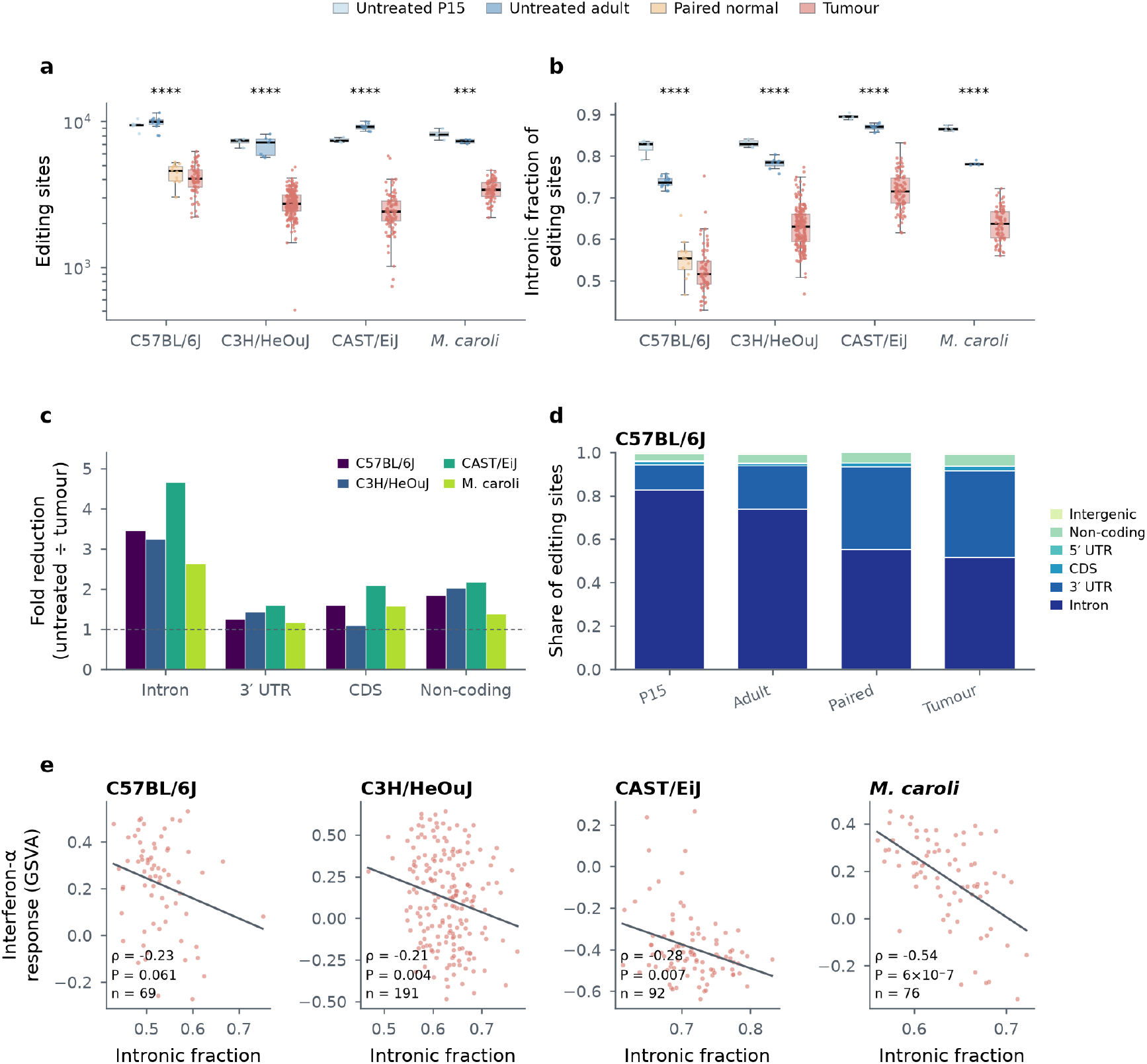
A-to-I editing is reduced in exposed tissue, selectively at introns. **a**, Total editing sites, the site counts behind the fold reductions in the text, shown beside **b** because count and fraction behave differently. **b**, Intronic fraction of editing sites by condition and background. A ratio, and therefore more robust to library depth than the raw site count. **c**, Fold reduction from untreated adult liver to tumour by region class. Introns lose most in every background; 3′ UTRs are nearly spared. Restricted to classes with enough sites for the ratio to be stable; 5′ UTR and intergenic sites number in the tens and are excluded. **d**, Share of sites by region and condition in C57BL/6J: a redistribution away from introns rather than a uniform loss. **e**, Intronic fraction against the hallmark interferon-α response in tumours, by background. Spearman with least-squares line.

The reduction is not uniform across the transcript (Fig. 5b,c). Partitioning sites by region, introns lose the most in every background (3.5-fold in C57BL/6J, 3.3 in C3H/HeOuJ, 4.7 in CAST/EiJ and 2.6 in *M. caroli*), while 3′ UTRs, the other major class, are nearly spared at 1.2- to 1.6-fold. Coding sites fall 1.1- to 2.1-fold. Because every class is counted in the same libraries, their relative composition is substantially less depth-sensitive than raw site counts, so a reduction concentrated in one class is unlikely to be explained by library depth alone. The consequence is a redistribution rather than a uniform loss (Fig. 5b,d). In C57BL/6J, taking median site counts per region, the intronic share of editing falls from 0.731 to 0.519 while the 3′ UTR share rises from 0.202 to 0.397 (the per-sample intronic fraction, used from here on, falls from 0.736 to 0.516), and the same reversal occurs in all four backgrounds. Editing is not simply reduced; it retreats from introns toward mature-transcript sequence.

Paired normals show the same region-specific pattern, at near-tumour magnitude. Intronic sites fall from 7,239 in untreated adult liver to 2,487, against 2,100 in tumours, while 3′ UTR sites are essentially unchanged (2,002 to 1,602, against 1,607 in tumours). The selectivity is therefore established before transformation, not acquired with it. One reading connects this to the processing-window argument set out in the Introduction. Most editing sites here are repeat-derived (∼87% in C57BL/6J), and repeat-derived duplexes that fall in introns are available to ADAR only while the intron is still present in the nascent transcript, whereas 3′ UTR duplexes persist in the mature message. A shortened window for co-transcriptional processing would therefore deplete intronic editing preferentially and leave 3′ UTR editing largely intact, which is the pattern observed. This is a consistency argument, not a demonstration: the data establish the selectivity, not its cause.

Editing level and circRNA abundance move inversely across conditions, which is the direction expected from ADAR activity on the duplex structures that promote backsplicing^37,40^. This association is pooled across conditions and is reported as descriptive: editing falls and circRNA abundance rises between normal liver and tumour, so a pooled correlation is produced by that difference whether or not the two are related within a state. For the same reason we do not use it to estimate how much of the circRNA enrichment is attributable to editing loss. To identify differential editing in hyper-edited regions, we defined hyper-edited clusters as regions with multiple RES interspersed by <10 bp. For each cluster we performed non-parametric analysis of editing-site density between conditions, identifying 544 (C57BL/6J), 223 (C3H/HeOuJ), 705 (CAST/EiJ) and 321 (*M. caroli*) differentially edited clusters after multiple-testing correction, with median absolute density differences of 0.137, 0.174, 0.175 and 0.200 respectively. Differential editing was detected at 1,184, 852, 1,072 and 921 sites, distributed across 184, 134, 152 and 136 genes. Per sample, a median of 553 clusters were defined in C57BL/6J (range 0–1,172, n = 104 samples), 373 in C3H/HeOuJ, 300 in CAST/EiJ and 405 in *M. caroli*. Differentially edited clusters span 3–430 bp with a median width of 28 bp in C57BL/6J, and comparable ranges in the other backgrounds (C3H/HeOuJ 3–350, CAST/EiJ 3–396, *M. caroli* 3–239 bp). Figure 6 shows editing sites across the *Spag16* locus, which carries one of the densest hyper-edited regions in C57BL/6J, expressed as the fraction of samples carrying each site so that groups differing in size remain comparable.

**Figure 6.**
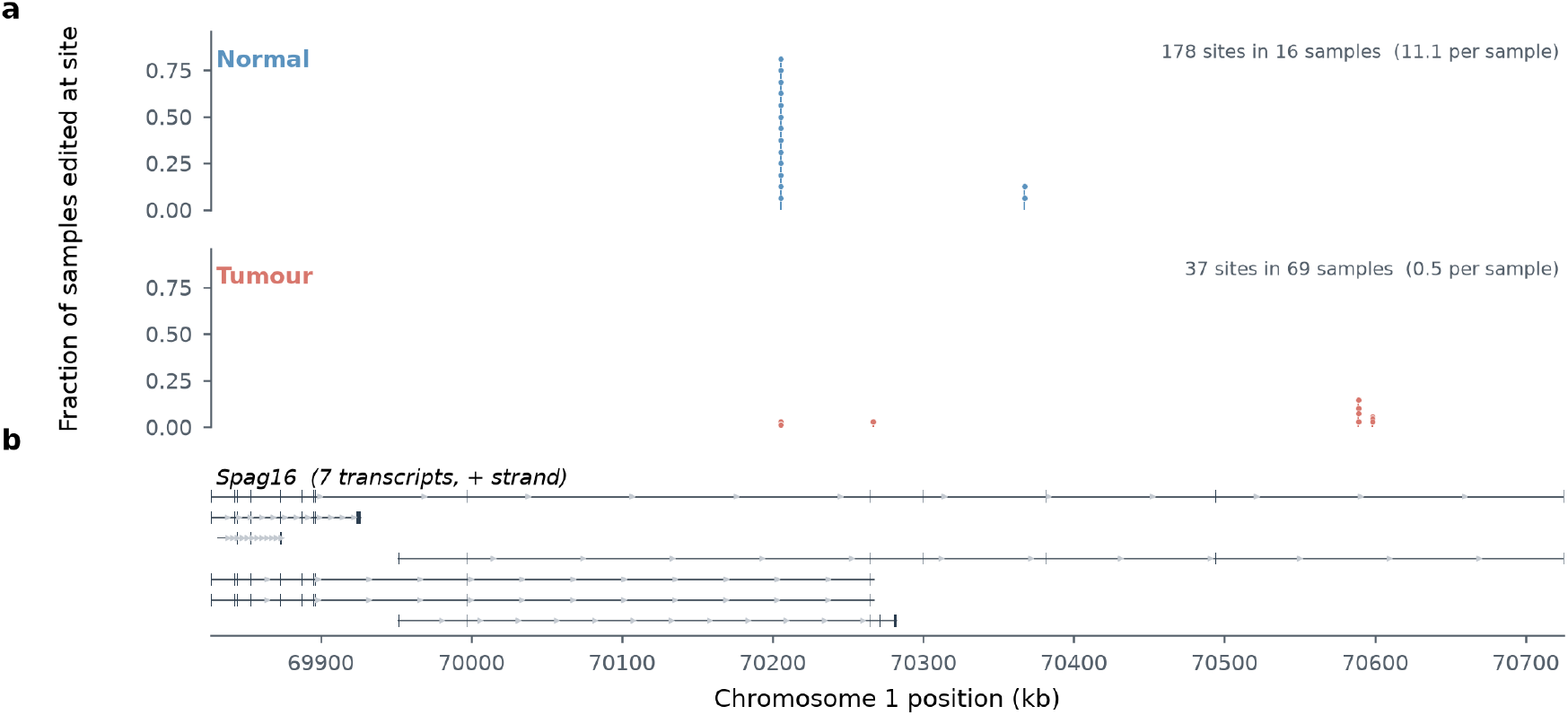
Editing sites across a hyper-edited locus, *Spag16* (C57BL/6J). **a**, Editing sites across *Spag16* in normal and tumour samples, plotted as the fraction of samples carrying each site so the two groups are comparable despite differing in size. **b**, Gene model, all annotated transcripts, exons as blocks.

Hierarchical clustering of editing levels shows differences between P15 and adult mice and between conditions. There are sub-clusters of tumour samples grouped by driver gene; however, the clustering of paired normal samples, which lack recurrent driver mutations, with tumour samples indicates similar editing profiles prior to transformation (Supplementary Fig. 5b). Editing site counts are negatively associated with cellularity in lesions of *M. caroli* (ρ = −0.456, *P* = 3.5 × 10^−5^, n = 76) and marginally so in C57BL/6J (ρ = −0.230, *P* = 0.058, n = 69; Supplementary Fig. 6), but the association does not survive on the depth-robust intronic fraction, where it is null in three backgrounds and reverses in C3H/HeOuJ (ρ = +0.316, *P* = 6.5 × 10^−6^).

Cellularity here is derived from ploidy and allele fraction (Methods), so it indexes how tumour-like a sample is, and conditioning on it would remove the effect under study. We found no association between editing and latency in any background on either measure (C57BL/6J ρ = +0.075, *P* = 0.55 on site counts; ρ = +0.163, *P* = 0.19 on the intronic fraction).

### *Adarb1* autoregulatory isoform usage is altered before transformation

Profiling *Adar* and *Adarb1* expression, we observed significant reductions in *Adarb1* that correlate with overall editing levels across strains. Gene-level expression does not by itself separate the two deaminases: correlating per-sample expression of each against editing partitioned by region class in C57BL/6J, intronic editing tracks *Adar* and *Adarb1* to the same degree (ρ = 0.584 for both, *P* = 2.7 × 10⁻¹⁰ and 2.8 × 10⁻¹⁰, n = 98), and the same holds at 3′ UTRs (ρ = 0.543 and 0.445) and coding sites (0.229 and 0.278). Both enzymes fall with transformation and a correlation on level alone cannot assign the loss to either. What is specific to *Adarb1* is not its level but its regulation.

*Adarb1* protein levels are normally regulated by auto-editing of its own locus, which generates isoforms producing non-functional proteins and non-coding RNAs^77^. Partitioning *Adarb1* transcript output by class shows that this regulation is altered, in the same direction in every background: retained-intron transcripts fall from about a fifth of output to a tenth (Cliff’s δ = 0.61 to 0.88, q = 6.8 × 10^−6^ to 0.018) and nonsense-mediated decay targets rise, significantly in C57BL/6J and CAST/EiJ (Supplementary Fig. 7). In C57BL/6J each change is already complete in paired normal liver and none progresses further with transformation. Because that autoregulatory loop is itself editing-dependent, its disruption is as readily a consequence of the editing loss as a cause of it.

### The post-transcriptional state precedes transformation

Twelve histologically normal liver samples were harvested from DEN-treated, tumour-bearing animals. These samples are the informative comparison in this study: they share the exposure of the tumours and the morphology of the controls. Four lines of evidence indicate that they carry the core tumour-associated post-transcriptional state without having transformed (Fig. 7).

**Figure 7.**
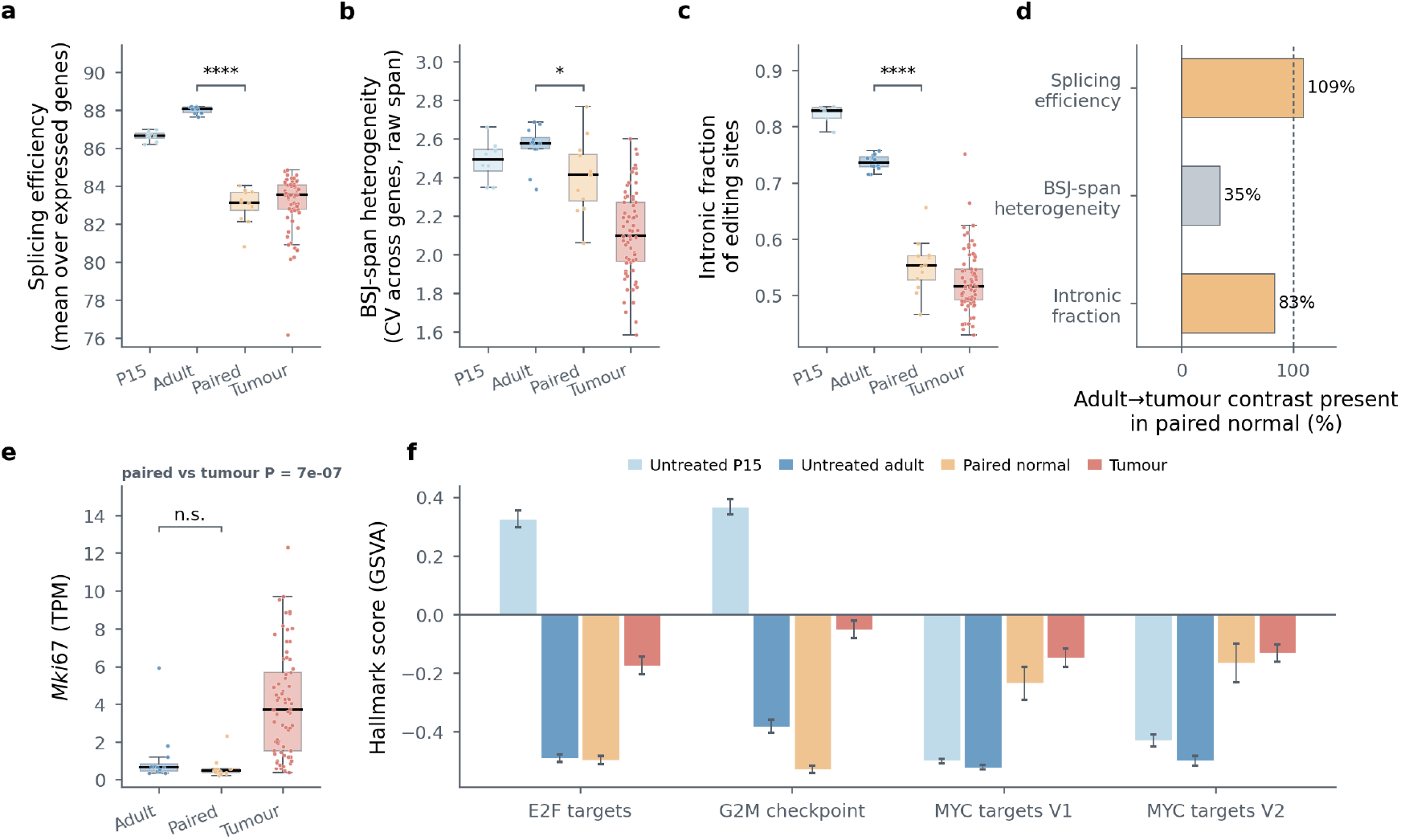
The post-transcriptional state is present before transformation. C57BL/6J. **a-c**, Splicing efficiency, cross-gene dispersion of maximal raw backsplice span, and the intronic fraction of editing sites, by condition. The circRNA quantity is the coefficient of variation, across genes, of each gene’s longest observed backsplice junction within a sample; it is a measure of how unevenly backsplice reach is distributed, not of its level, and is distinct from the per-sample mean relative span used in Fig. 8. Boxes are quartiles with the median; points are samples. Brackets compare paired normals with untreated adult liver. Splicing efficiency and editing place paired normals at or near the tumour value; dispersion falls from untreated adult liver through paired normal to tumour, with paired normal already differing from untreated adult (*P* = 0.019). **d**, Share of the untreated-to-tumour contrast already present in paired normals, by measure: 109% for splicing efficiency, 35% for backsplice-span dispersion and 83% for intronic editing. Splicing and editing are therefore at or near their lesion-state values before morphological transformation, while backsplice architecture has begun to reorganise without being fully shifted. **e**, Mki67 expression by condition. Paired normals are indistinguishable from untreated adult liver (*P* = 0.19) and far below tumours (*P* = 7.3 × 10⁻⁷), Mann–Whitney. **f**, Hallmark proliferation programmes by condition, median with standard error. Cell-cycle sets place paired normals at or below untreated adult liver; MYC targets, which are dominated by biosynthetic and RNA-processing genes, are already at tumour level. Occult tumour content would raise both, and does not.

#### They are transcriptionally distinct from untreated liver

Differential expression between paired normal and untreated adult liver identifies 7,895 protein-coding genes (adjusted *P* < 0.05), within the range of the dysplastic contrasts in the same background, from 7,520 (*Egfr*-driven) to 10,505 (*Braf*-driven). Exposure therefore produces a widespread transcriptional response in histologically normal tissue, comparable in breadth to the lesion contrasts in this background.

#### They already carry most components of the tumour-associated post-transcriptional state (Fig. 7a–d)

Splicing efficiency and intronic editing are at or near tumour values, circRNA abundance is tumour-like (Fig. 4c,f), and cross-gene heterogeneity of absolute backsplice span has already shifted in the tumour direction. circRNA abundance profiles in paired normal samples are not obviously different from tumour samples, where both differ from untreated P15 and adult liver. Intronic RES reductions are likewise observed in paired normal samples, and paired normals cluster with tumours on editing profiles despite lacking recurrent driver mutations.

#### They have not clonally expanded

Mutation burden in paired normal samples has a median of 337 single-nucleotide variants (range 66–3,956, n = 12) against a tumour median of 78,843, a 234-fold difference (*P* = 5.2 × 10^−8^), and these samples are devoid of recurrent driver mutations. The low callable burden reflects polyclonality rather than absence of damage: in polyclonal tissue each mutation sits at low variant allele fraction and is individually uncallable, so a callable burden is a readout of clonal expansion rather than of exposure.

#### They are not cycling (Fig. 7e)

*Mki67* expression in paired normal samples is indistinguishable from untreated normal liver (*P* = 0.19) and significantly lower than in tumours (*P* = 7.3 × 10^−7^). Because a single marker gene is a limited basis for this inference, we tested it against multi-gene programmes scored per sample by gene set variation analysis. Cell-cycle programmes agree (Fig. 7f): paired normals match untreated adult liver for E2F targets (median −0.497 against −0.482, *P* = 0.34) and sit *below* it for the G2M checkpoint (−0.531 against −0.371, *P* = 7.7 × 10^−5^), while tumours exceed them on both (*P* = 8 × 10^−8^ and 6 × 10^−8^). MYC targets behave differently, and informatively. Paired normals are elevated relative to untreated liver (−0.234 against −0.519, *P* = 3.7 × 10^−5^ for MYC_TARGETS_V1; −0.166 against −0.473, *P* = 0.002 for V2) and are indistinguishable from tumours (*P* = 0.18 and 0.22). MYC target sets are dominated by biosynthetic and RNA-processing genes rather than cell-cycle machinery, so the pattern is a growth and biosynthesis programme already engaged in tissue that is not dividing. The alternative reading, that these samples contain occult tumour tissue, is not supported by the cell-cycle programmes, which would be elevated alongside MYC targets under contamination and are not.

The post-transcriptional state is therefore present in tissue that is neither clonal nor cycling. Because the number of editing sites detected scales with library depth, the informative quantity is the *proportion* of edited sites that are intronic, which is a ratio and therefore more robust to library depth than the raw site count. It falls monotonically with exposure and transformation in C57BL/6J (Fig. 7c), 0.828 in untreated P15 liver, 0.736 in untreated adult liver, 0.553 in paired normals and 0.516 in tumours. Paired normals sit much closer to the tumour value than to the untreated one, and median site counts behave the same way (4,559 against 4,048 in tumours and 9,898 in untreated adults). The reduction in intronic editing that accompanies transformation is thus largely established in tissue that has not transformed, with 83% of the untreated-to-tumour contrast already present (Fig. 7d).

The C57BL/6J animals providing paired normals were sampled 34–37 weeks after a single juvenile dose (median 35); each histologically normal sample was harvested from the same animal, and at the same time, as one of the sequenced tumours, so the interval is that animal’s own. This establishes that the state is durable and can be present in exposed tissue without local clonal transformation. The same ordering holds in the other three backgrounds for the groups available there, with intronic fractions of 0.78–0.87 in untreated adult liver against 0.63–0.71 in tumours (Supplementary Data 6).

#### Core components of the state do not deepen with progression

Comparing groups two at a time cannot distinguish a change that is complete before transformation from one this cohort is merely underpowered to follow, so we placed the three measures on the ordered series this background supports: paired normal liver (n = 12), dysplastic nodule (n = 58) and hepatocellular carcinoma (n = 11). None of the three moves along it. Splicing efficiency reads 83.1, 83.6 and 83.0 across the three stages, the intronic fraction of editing 0.553, 0.513 and 0.546, and per-sample mean relative backsplice span 0.0283, 0.0250 and 0.0296; the trend coefficient lies between −0.05 and −0.02 for all three and no test approaches significance (Spearman against ordinal stage, *q* > 0.78 after correction). Set against untreated adult liver at 88.1, 0.736 and 0.0277, the change in splicing efficiency and editing has occurred by the paired-normal stage and no additional change is detected with progression. The third of these is the per-sample mean, which is flat in every contrast, and not the cross-gene span dispersion of Figure 7b, which behaves differently.

Backsplice architecture behaves differently and qualifies the section above (Fig. 7b,d). Its per-sample mean relative span shows no detectable shift: paired normals do not differ from untreated liver (0.0283 against 0.0277), nor do tumours (0.0250, *P* = 0.38). What moves is the distribution of span across genes. The cross-gene dispersion of each gene’s maximal raw backsplice span falls from 2.58 in untreated adult liver to 2.41 in paired normals and 2.10 in tumours (*P* = 0.019 and 3.0 × 10^−7^), so the reorganisation has begun before transformation, with 35% of the adult-to-tumour contrast already present against 109% for splicing efficiency and 83% for editing (Fig. 7d). The adult-to-tumour reduction reproduces in C3H/HeOuJ and CAST/EiJ and not in *M. caroli* (Supplementary Fig. 4); the pre-transformation timing can be resolved only in C57BL/6J.

The series itself detects change where change occurs. Scoring the same libraries for immune and stromal composition, several populations do move monotonically along the same series, monocytes from 7.23 through 5.19 to 4.70 (ρ = −0.54, *q* = 4 × 10^−6^), endothelial cells, eosinophils and basophils in the opposite direction (ρ = +0.39 to +0.43, all *q* ≤ 0.001), and the series resolves coefficients as small as 0.26. The comparison also separates the two kinds of marker within one modality: macrophages rise sharply between untreated and paired normal liver, 5.46 to 7.18, and are then flat across dysplasia and carcinoma (ρ = −0.05), behaving as the post-transcriptional measures do, while monocytes and eosinophils track the lesion. Exposure and progression leave distinguishable signatures in the same samples, and the core post-transcriptional state is established before lesion progression rather than deepening with it.

The comparison can also be made within animals. Each of the twelve C57BL/6J animals providing a paired normal also provides five or six lesions, carrying two to four different driver mutations. Placing splicing efficiency, per-sample mean relative backsplice span and the intronic fraction of editing on a common scale, each animal’s paired normal lies a median of 1.3 standard deviations from its own lesions and 3.1 from untreated adult liver, and the lesions are the closer of the two in every animal taken separately. Within an animal, lesions of different drivers are equidistant from the paired normal (permutation *P* = 0.85 across the five single-driver classes; *Braf* against *Hras* in the six animals carrying both, *P* = 1.0), and a paired normal is no closer to its own animal’s lesions than to those of other animals (*P* = 0.97). Because drivers differ in latency^5^, lesions of different drivers within an animal differ in age, so this is a further indication, independent of the dysplasia-to-carcinoma series, that the state does not deepen once transformation has occurred (Supplementary Data 1).

#### One splice measure does continue to change with pathological progression

In C57BL/6J the novel splice proportion rises from a median of 19.65% in dysplastic nodules (n = 58) to 21.47% in hepatocellular carcinomas (n = 11, *P* = 0.0025), and the difference persists when animals rather than lesions are the resampling unit (+1.82 percentage points, 95% CI +0.78 to +5.07). It is not attributable to tumour content: within C57BL/6J tumours NSP is uncorrelated with genomic cellularity (ρ = +0.021, *P* = 0.87). The other backgrounds contain two to four carcinomas and so provide no informative replication. Because NSP records the repertoire of tumour-specific junctions rather than their abundance per cell, we read this as continued expansion of detectable cryptic splice-site use after canonical splicing efficiency has reached its altered state, rather than as progressive worsening of the splicing defect itself.

#### The principal lesion-level results are robust to within-animal dependence

Because lesions from the same animal are not independent, each principal contrast was repeated with animals, rather than lesions, resampled with replacement. Resampling this way preserves the adult-to-tumour reductions in splicing efficiency and intronic editing in all four backgrounds, and the reduction in cross-gene backsplice-span dispersion in C57BL/6J, C3H/HeOuJ and CAST/EiJ but not *M. caroli*. The C57BL/6J paired-normal contrasts remain resolved, including the smaller pre-transformation change in span dispersion (95% CI −0.336 to −0.037). Conversely, the absence of further progression from dysplasia to carcinoma in splicing efficiency, intronic editing and span dispersion is retained under the same procedure, with every interval spanning zero. The positive pre-transformation contrasts therefore survive cluster-aware inference, while no additional change from dysplasia to carcinoma is detected in these measures under the same procedure (Supplementary Data 1).

We asked whether the depth of this state scales with the mutagenic dose each animal received. Within the paired normal samples, exposure to the first of the two DEN mutational signature components (den1sig and den2sig; Methods) spans roughly fifty-fold (0.015–0.775) while the outcome is held at zero, which makes them the only group in which exposure varies independently of transformation. We correlated per-sample splicing efficiency, circRNA abundance, expression and per-sample mean relative backsplice span against den1sig and den2sig, with two-sided permutation *P* values.

#### No dose-response was detected

in paired normals (n = 12; all |ρ| ≤ 0.48, all *P* ≥ 0.114, where |ρ| ≥ 0.59 is required for significance). Two things make that null unsurprising. The response has largely already happened: paired normals have completed the whole untreated-to-tumour change in splicing efficiency (88.1 to 83.1, tumour median 83.5) and 83% of it in intronic editing fraction (0.736 to 0.553, tumour 0.516), so a gradient is being sought at the plateau. And the predictor is poorly estimated here, resting on a median of 337 mutations against 76,675 in nodules.

Testing the gradient where it can be tested, within dysplastic nodules, gives no consistent dose-response either. Splicing efficiency falls with den1sig and mutation burden in *M. caroli* (ρ = −0.295 and −0.309, cluster-bootstrap CIs excluding zero) but in no other background. Intronic editing fraction rises with den1sig, den2sig and mutation burden in C3H/HeOuJ (ρ = +0.22 to +0.32), opposite in sign to the between-group relationship and likely reflecting the cellularity association described above. Span shows no association anywhere. Within the exposure range sampled we therefore detect no consistent graded relationship between these measures and the exposure estimates.

### The state has an animal-level component

Eighty-nine animals contribute more than one dysplastic nodule with a transcriptome. Because those nodules share a germline, a dose, an age at treatment and at sacrifice, and a cage, comparing them separates what differs between lesions from everything that differs between animals. We quantified the fraction of variance lying between animals rather than within them, as a one-way random-effects intraclass correlation, restricted to dysplastic nodules so that lesion type is held constant, and tested against a null obtained by permuting animal labels. The between-animal share of variance is given in Table 1. The per-sample summaries used here and in the coordination analysis are defined in Methods.

**Table 1.** Between-animal share of variance in each measure, by background. Intraclass correlation across dysplastic nodules from animals contributing at least two lesions. Bold marks estimates whose permutation *P* is below 0.05.

| background | animals | nodules | splicing efficiency | mean relative span | editing |
| --- | --- | --- | --- | --- | --- |
| C3H/HeOuJ | 39 | 161 | **0.39** ($< 2 \times 10^{-4}$) | **0.21** (0.001) | **0.35** ($< 2 \times 10^{-4}$) |
| *M. caroli* | 19 | 46 | -0.06 (0.63) | 0.32 (0.050) | **0.43** (0.008) |
| C57BL/6J | 12 | 55 | -0.10 (0.83) | -0.04 (0.60) | -0.01 (0.49) |
| CAST/EiJ | 19 | 46 | -0.05 (0.59) | 0.04 (0.39) | -0.19 (0.85) |

In C3H/HeOuJ, the background contributing most animals, between a fifth and two fifths of the variance in all three measures is shared between nodules of the same animal. *M. caroli* agrees for editing, with mean relative span at the threshold (*P* = 0.050). A substantial part of the post-transcriptional state is therefore a property of the animal rather than of the individual lesion, consistent with an animal-level field component, though the intraclass correlation cannot distinguish field biology from other factors shared within an animal.

The two backgrounds returning null estimates have the two narrowest sampling windows, C57BL/6J 34–37 weeks and CAST/EiJ 36–43, against 22–39 for C3H/HeOuJ and 56–80 for *M. caroli*; C57BL/6J also contributes the fewest animals. An intraclass correlation requires between-animal variance to exist before it can be detected. The null estimates therefore do not exclude an animal-level component, particularly where the sampled between-animal range is narrow.

### Lesion genotype and ploidy

The C57BL/6J cohort carries *Ctcf;AlbCre* HE;HE (36 lesions) and HE;WT (33) in near-equal numbers, with the Cre allele driving hepatocyte-specific deletion. Because CTCF organises chromatin loops and influences elongation, the account above predicts an effect on splicing efficiency and backsplice span; and because differential expression between these genotypes returns only thirteen genes, any post-transcriptional shift would be a clean dissociation. There is none. Splicing efficiency is 83.5% in both genotypes (Cliff’s δ = −0.005, *P* = 0.98), per-sample mean relative backsplice span 0.0253 against 0.0250 (δ = −0.09, *P* = 0.54) and the intronic fraction of editing 0.514 against 0.516 (δ = −0.05, *P* = 0.71); cellularity is balanced between genotypes (*P* = 0.45). The absence of a detectable Ctcf-associated effect does not identify the route by which elongation may be altered; intragenic methylation remains one candidate.

#### Editing is elevated in whole-genome-duplicated lesions

*M. caroli* carries 29 duplicated lesions against 44 clonal. Editing is higher in the duplicated group (δ = 0.45, *P* = 0.0014), while splicing efficiency and per-sample mean relative backsplice span are unchanged (*P* = 0.32 and 0.89). This comparison is not adjusted for cellularity, although the groups differ markedly (0.349 against 0.640, *P* = 7 × 10⁻¹³). The reason is definitional rather than statistical. Cellularity in this cohort is not measured: it is computed as ploidy × the modal variant allele fraction, with ploidy set to two for every sample except *M. caroli* tumours showing no mutational asymmetry, which are inferred to be whole-genome duplicated and assigned ploidy four^45^. The duplicated group in this contrast is precisely the set of samples to which that assignment was applied. The covariate therefore embeds the exposure, and conditioning on it conditions on a re-expression of the variable under test, which is exactly what is observed: adjustment removes the association entirely (*P* = 0.39).

Two further quantities behave as that construction predicts. The fraction of segments showing allelic imbalance falls to zero in the duplicated group, which follows by definition, since absence of mutational asymmetry is the criterion by which duplication was called. VAF skew inverts in sign between the groups (+0.31 against −0.30), consistent with the shifted allele-fraction distribution a doubled genome produces. The one quantity measured rather than derived shows no difference: the lesions are the same size (diameter 4.55 against 4.41 mm, *P* = 0.95). We therefore report the contrast unadjusted.

That editing moves while splicing efficiency and span do not argues against a kinetic explanation for this particular contrast, since a broad kinetic change might be expected to perturb more than editing alone. A simpler reading is that a doubled genome raises the abundance of the double-stranded substrates ADAR acts on. We cannot separate this from damage burden: duplicated lesions also carry more mutations (73,398 against 49,973, *P* = 0.0015), and either could account for the difference. Separating them requires an estimate of tumour cell content that is orthogonal to allele fractions and ploidy. No such estimate exists in this cohort: the lesion descriptors record diameter and the histological scores, but their cellularity field is the genomic quantity defined above, not a pathologist’s assessment. An image-derived measure, nuclear-size dispersion from the *M. caroli* whole-slide images (S-BSST384), would supply one, and is deferred to the histological companion study.

### Editing loss and the immune compartment

To characterise non-hepatocyte composition we profiled tissue-specific markers in each RNA-seq sample using mMCP-counter^78^. Basophil and eosinophil scores are higher in tumours (Supplementary Fig. 8). Loss of A-to-I editing has consequences beyond transcript diversity. Unedited double-stranded RNA, particularly duplexes formed by inverted repeat elements, which constitute the majority of the sites identified here, is the ligand for the cytosolic sensors MDA5, PKR and ZBP1, and editing capacity is a determinant of whether such transcripts activate innate immune signalling^79^. A reduction in editing concentrated at repeat-derived intronic sites therefore makes a specific prediction: samples that have lost the most intronic editing should show the strongest interferon response.

They do. Scoring each sample against the MSigDB hallmark collection and correlating within condition, the intronic fraction of editing is negatively associated with the interferon-α response in tumours of every background, ρ = −0.227 (*P* = 0.06) in C57BL/6J, −0.207 (*P* = 0.004) in C3H/HeOuJ, −0.277 (*P* = 0.007) in CAST/EiJ and −0.536 (*P* = 5.9 × 10^−7^) in *M. caroli* (Fig. 5e). The interferon-γ response follows the same direction, reaching significance in CAST/EiJ (−0.206) and *M. caroli* (−0.416). Paired normals show the same sign, at twelve samples, where |ρ| ≥ 0.58 reaches significance.

The specificity of this association is what makes it interpretable. If the relationship simply tracked immune infiltrate, generic inflammatory programmes should move with it. They do not: the inflammatory response and IL6–JAK–STAT3 signatures lie between −0.01 and −0.12 across all four backgrounds and reach significance nowhere. The association is confined to the interferon axis, which is the arm of innate immunity that cytosolic double-stranded RNA sensing activates. This is consistent with unedited duplex RNA acting as ligand, and is not what non-specific immune infiltration would produce.

The measure matters here. Raw counts of edited sites scale with library depth, and depth also raises the resolution of expression-based scores; correlating raw counts inverts the sign of every association above. The depth-robust intronic fraction is therefore the more interpretable quantity here, and it is the quantity reported.

mMCP-counter deconvolution (Supplementary Fig. 8) shows shifts in basophil, eosinophil, macrophage and monocyte populations between conditions. This is an association, not a demonstrated mechanism. Editing loss and interferon activation could both follow from transformation without either causing the other, and nothing here establishes direction. The data argue against the simplest alternative, that the relationship is merely a by-product of immune cell content. That argument does not rest only on the specificity of the signature: correlating the interferon score against each of the fourteen immune and stromal populations estimated by mMCP-counter within C57BL/6J tumours returns no association surviving correction across populations (n = 69; smallest *q* = 0.25, eosinophils). The interferon response in these samples is not accounted for by how much immune tissue they contain. Editing itself is not independent of composition in the same way: across the same fourteen populations, the intronic fraction of editing is associated with basophil (ρ = −0.39), eosinophil (ρ = +0.37) and memory B cell (ρ = +0.36) scores (each *q* = 0.010, n = 69), so part of the variation in editing is shared with infiltrate even though the interferon association is not.

### The editing loss does not require the mutagen

Ten C3H/HeOuJ lesions in this cohort arose **spontaneously**, without DEN. They provide a control that no dose or timing analysis can: tissue that transformed without the exposure. Their intronic editing fraction is 0.543, against 0.630 in DEN-induced C3H/HeOuJ tumours and 0.785 in untreated adult liver of the same background. The loss is therefore not a signature of the mutagen. If anything it is deeper without it, though ten samples will not support a strong claim about the difference.

This bears directly on how the state should be described. A change seen only after chemical exposure invites reading as a lesion-specific damage response; the same change in spontaneous lesions of the same background makes that reading untenable and points instead to something general about the transformed, or transforming, state. It also constrains the interpretation of *The post-transcriptional state precedes transformation*: the paired normals establish that the change precedes transformation, and the spontaneous lesions establish that the editing loss does not require DEN exposure. Together they indicate that the state accompanies tissue that is transforming, or at risk of doing so, rather than being a direct signature of DEN.

### Coordination between processes, and its breakdown

The three changes described above are usually reported as separate phenotypes. We asked whether they are manifestations of a common state, which is a question about shared structure rather than about how much of gene expression each explains.

Measure selection matters here and constrains what can be concluded. Several quantities in this study are not independent of one another: the circRNA abundance proportion is BSJ/(BSJ+CSJ), so it shares its denominator with the numerator of the splicing efficiency index and rises mechanically whenever canonical splicing falls, and the novel splice proportion is a further ratio over the same junction reads. Shared structure across such measures would be partly definitional. We therefore selected three quantities that are independent by construction: splicing efficiency (a read ratio at annotated junctions), per-sample mean relative backsplice span (a genomic coordinate distance normalised to gene length, not a read count) and RNA editing level (nucleotide mismatches). Gene expression is excluded from this analysis because it is downstream of all three and would absorb their shared variance; it is the outcome of the apportionment specified below, not an input here.

At the level of individual features, no factor is shared: a joint decomposition across the three matrices yields a leading factor explaining 21% of splicing variance and ∼2% of the others. Feature-level variance is dominated by gene-specific noise, and the claim concerns sample-level state.

At the level of per-sample summaries the structure is legible, and it separates three states (Table 2; Fig. 8a,b). Alongside untreated controls and tumours we include the paired normal samples, which share the tumours’ exposure and differ only in not having transformed.

**Figure 8.**
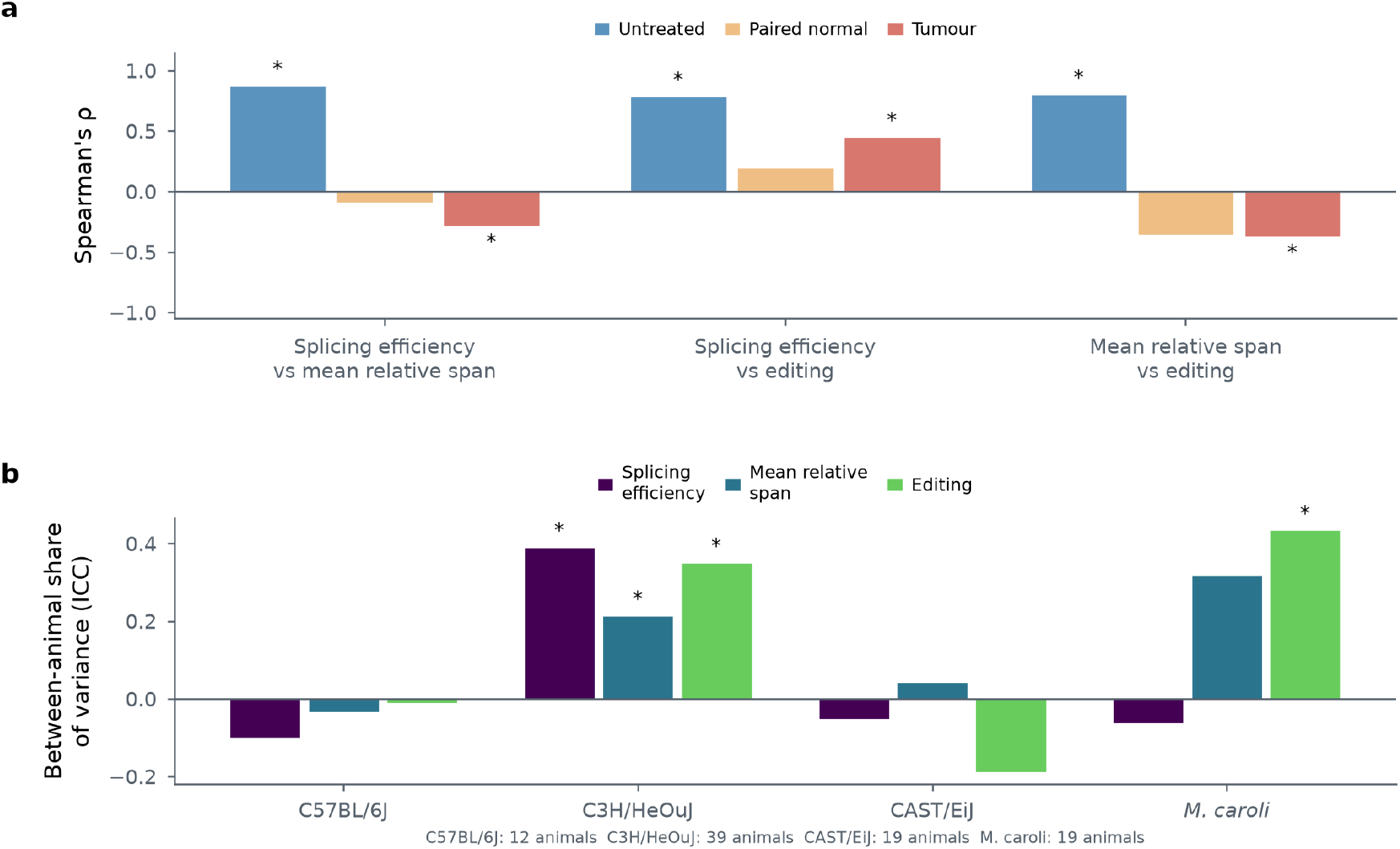
Coordination between post-transcriptional processes is lost on exposure, and a substantial share of the state belongs to the animal. **a,** Pairwise Spearman correlations between per-sample splicing efficiency, per-sample mean relative backsplice span and editing level, within condition, in C57BL/6J. In untreated liver all three co-vary positively (ρ +0.78 to +0.87). That structure is absent in exposed tissue; paired normal liver is indistinguishable from tumour (Fisher’s z, *P* = 0.56 and 0.95) while both differ from untreated liver (*P* = 0.003 and 0.002). Pooling conditions obscures this, because the estimates have opposing signs and cancel. Asterisks mark correlations significant at *P* < 0.05. **b,** Intraclass correlation for the same three measures across dysplastic nodules from the same animal, by background, as the between-animal share of total variance; restricted to dysplastic nodules so that lesion type is held constant. Asterisks mark estimates exceeding a null obtained by permuting animal labels. In C3H/HeOuJ (39 animals, 161 nodules) between a fifth and two fifths of the variance in all three measures is shared within animals. C57BL/6J and CAST/EiJ return null estimates and have the two narrowest sampling windows; an intraclass correlation requires between-animal variance to be present before it can be detected.

**Table 2.** Coordination between measures across three states, C57BL/6J. Spearman correlations within condition, with the two-sided *P* in parentheses. Bold marks correlations significant at 0.05.

| relationship | untreated (n = 11) | paired normal (n = 12) | tumour (n = 69) |
| --- | --- | --- | --- |
| splicing efficiency vs editing | **+0.78 (0.004)** | +0.20 (0.54) | **+0.45 ($1 \times 10^{-4}$)** |
| splicing efficiency vs mean relative span | **+0.87 ($5 \times 10^{-4}$)** | -0.09 (0.78) | **-0.29 (0.016)** |
| mean relative span vs editing | **+0.80 (0.003)** | -0.36 (0.26) | **-0.38 (0.002)** |

In untreated liver the three processes co-vary positively: samples with more efficient canonical splicing also show higher editing and longer mean relative backsplice spans. That three-way positive coordination is absent in exposed tissue, whether or not it has transformed.

Testing the differences directly (Fisher’s z), untreated liver differs from both other states on the two relationships involving mean relative span, against paired normals, *z* = 2.95 (*P* = 0.003) and *z* = 3.03 (*P* = 0.002); against tumours, *z* = 4.38 (*P* = 1 × 10^−5^) and *z* = 3.99 (*P* = 7 × 10^−5^), while paired normals and tumours do not differ from one another (*P* = 0.56 and 0.95). The splicing–editing coupling shows no significant difference between any pair of states. Three points follow. First, the decoupling of mean relative span from the splicing–editing axis is already present in exposed, histologically normal tissue and precedes local morphological transformation: tissue that is non-proliferating and lacks tumour-like clonal expansion or recurrent drivers has already lost it and is indistinguishable from tumour in this respect. Second, this is invisible when conditions are pooled (ρ = −0.14 and −0.19, neither significant), because the untreated and exposed groups have opposing signs that cancel; analyses of this kind must be conducted within stratum. Third, exposure changes individual processing outputs as well as the structure relating them, and that structural change, like the level changes, precedes any morphological evidence of transformation.

The three-state comparison is available only in C57BL/6J. Paired normal liver was harvested from twelve C57BL/6J animals, two C3H animals and none of the *M. m. castaneus* or *M. caroli* cohorts, so the contrast that dates the loss of coordination cannot be repeated in another background and we do not present the timing claim as a cross-background result.

The two-state form of the comparison, untreated liver against lesions, is available in all four, and the same directional change recurs where it can be estimated (Table 3). Both relationships involving mean relative span are positive in untreated liver and fall to near zero or negative in lesions in every background where the estimate is defined, while the splicing–editing coupling, the one relationship that does not differ between states in C57BL/6J, stays positive in both states everywhere.

**Table 3.** Coordination between measures in untreated liver and in lesions, by background. Spearman correlations within each condition, with Fisher’s *z* for the difference between them. Untreated liver contributes 11, 6 and 7 libraries and lesions 69, 195 and 77 in C57BL/6J, C3H/HeOuJ and CAST/EiJ respectively; *M. caroli* has three untreated libraries with all three measures and is not estimated. Bold marks within-condition correlations significant at *P* < 0.05.

| background | relationship | untreated | lesion | $z$ | $P$ |
| --- | --- | --- | --- | --- | --- |
| C57BL/6J | splicing efficiency vs mean relative span | **+0.87** | **−0.29** | 4.38 | $1 \times 10^{-5}$ |
| C57BL/6J | mean relative span vs editing | **+0.80** | **−0.38** | 3.99 | $7 \times 10^{-5}$ |
| C57BL/6J | splicing efficiency vs editing | **+0.78** | **+0.45** | 1.52 | 0.13 |
| C3H/HeOuJ | splicing efficiency vs mean relative span | +0.71 | +0.10 | 1.37 | 0.17 |
| C3H/HeOuJ | mean relative span vs editing | +0.60 | −0.10 | 1.36 | 0.17 |
| C3H/HeOuJ | splicing efficiency vs editing | **+0.94** | **+0.61** | 1.80 | 0.072 |
| CAST/EiJ | splicing efficiency vs mean relative span | +0.61 | +0.20 | 0.98 | 0.33 |
| CAST/EiJ | mean relative span vs editing | **+0.79** | −0.14 | 2.33 | 0.020 |
| CAST/EiJ | splicing efficiency vs editing | +0.68 | **+0.53** | 0.47 | 0.64 |

The non-reference estimates are therefore directional rather than powered. Read together with the direction of each individual change, reduced splicing efficiency and editing together with the circRNA repertoire and architectural changes reported in the preceding sections, what recurs across backgrounds is the coordination structure and the direction in which it breaks; what remains specific to C57BL/6J is the timing.

## Discussion

### Coordination, and what transformation does to it

The most direct evidence that these processes are related is that they co-vary. In untreated C57BL/6J liver, splicing efficiency, per-sample mean relative backsplice span and editing level move together across samples, pairwise correlations of +0.78 to +0.87. That is not a statement about mean differences between groups; it is a statement that the three are, in normal tissue, expressions of a common state. In tumours the splicing–editing coupling survives while both relationships involving backsplice invert. Exposure and transformation therefore do two things: they change several of the individual processing outputs, and they dismantle part of the structure relating them. The second is invisible to any analysis that compares group means, and invisible to a pooled correlation, because the opposing signs cancel. Notably the mean relative-span summary itself shows no corresponding level shift.

### One kinetic shift, and where it stops explaining things

The changes reported here are usually studied separately, and each has its own literature: splicing efficiency, circRNA biogenesis, RNA editing. We find these phenotypes together in the same samples across four mouse backgrounds, following one exposure. A parsimonious hypothesis is that they reflect a shared alteration in post-transcriptional processing state. Acute polymerase stalling at transcription-blocking lesions is well-established^18,19,26,28^ and, on its own, would predict slower elongation and increased exon inclusion^49^. That is the acute response. What we measure, months later, is the tissue after repair. Gene-body hypomethylation^80^ is one candidate route to a changed elongation or processing regime, since it is associated with faster elongation and altered splice-site selection^29–31^. The two de novo DNA methyltransferases point the same way: *Dnmt3a* and *Dnmt3b* are expressed at lower levels in tumours than in untreated adult liver in every background (1.4-to 2.1-fold and 1.9-to 3.4-fold lower, respectively), and in C57BL/6J paired normal liver they are already as low as in tumours. We did not measure DNA methylation or elongation rate, so this is consistent with that route rather than evidence for it. A shortened window for co-transcriptional splice-site recognition predicts a falling canonical splicing efficiency and, because linear splicing now lags transcription, altered competition among the donor–acceptor configurations available for backsplicing. What we observe matches the second prediction in architectural rather than arithmetic form: tumours reach distal configurations rarely seen in normal liver and the cross-gene distribution of span is reorganised, while per-sample mean relative span does not rise. The reduction in SEI and the reorganisation of backsplice architecture are consistent with two readouts of the same underlying shift. Reduced splicing efficiency also has consequences for expression in its own right, since errors in RNA processing can trigger nonsense-mediated decay^81^ or leave transcripts retained at nuclear speckles before degradation^82,83^.

We note that positive relationships between elongation speed and circRNA formation have been reported^84^, and that a competing account exists in which pre-mRNA processing machinery becomes limiting rather than polymerase becoming faster^15,32^. The two are not mutually exclusive and are distinguishable: a capacity limitation predicts that the loss concentrates in highly expressed, intron-dense genes, whereas a kinetic account predicts dependence on gene length and intragenic methylation with weaker dependence on expression.

### The timing is the finding

Reduced *Polr2a* expression persisting months after treatment, contrary to reports of post-repair recovery^26,48^, was the first indication that these consequences outlast their cause. The paired normal samples establish it. Tissue that a pathologist classifies as normal, from an animal bearing tumours, already carries tumour-like splicing efficiency, circRNA abundance and editing while lacking their clonal mutation burden and showing no elevation of proliferation, and its backsplice architecture has begun to reorganise without being fully shifted. Placed on the ordered series that follows, from that tissue through dysplastic nodule to carcinoma, splicing efficiency, editing and mean relative span move no further, while immune and stromal populations scored on the same libraries move monotonically along it and the novel splice proportion continues to rise. The core state is therefore established before local morphological transformation and does not require it, although some components continue to evolve within lesions.

This distinction matters for how the result is read. Post-transcriptional dysregulation here marks the *field* rather than the *lesion*: it is present in exposed tissue whether or not that tissue has clonally expanded. We use “transformation” in the sense of the process by which exposed tissue becomes neoplastic, not as a synonym for clonal conversion, the paired normal samples are explicitly not the latter. Read this way, the claim is stronger than a tumour-versus-normal comparison, because it identifies molecular change that precedes anything morphologically identifiable.

### What mutation does not explain

Splice-region mutations are associated with the expression of under 0.6% of genes tested, and splice sites are mutated at no more than the intronic rate. The splicing phenotype is not written in the sequence of its substrate. Together with the presence of the phenotype in samples devoid of recurrent drivers, this argues that the post-transcriptional state is regulatory rather than a direct genetic consequence of the mutations defining the lesions.

### Editing loss is associated with interferon activation

The 2.2-to 3.8-fold depletion of A-to-I editing at repeat-derived intronic sites is reported here as a transcriptome phenomenon, but unedited inverted-repeat duplexes are the substrate for MDA5, PKR and ZBP1. Editing does covary with immune composition, tracking basophil, eosinophil and memory B cell scores in C57BL/6J tumours, but the interferon response it is associated with is not explained by immune content (Results).

### Cross-background conservation

The same directions of change in four mouse backgrounds that differ in latency, susceptibility and evolutionary distance is the strongest evidence that the core changes are not peculiar to one genetic background.

### Limitations

The coordination analysis and the timing claim that follows from it rest on C57BL/6J alone, because paired normal liver was collected from twelve C57BL/6J animals, two C3H animals and none of the other two backgrounds. That is a property of the cohort rather than of the analysis, and no reanalysis can recover it. The individual changes in splicing efficiency, circRNA abundance and repertoire, and editing recur across all four backgrounds, and the raw-span dispersion phenotype in three of them; the structure relating them, and when that structure is lost, are established in one.

These are also bulk measurements over tissue of mixed cellularity; we control for tumour cell content where possible but cannot resolve cell-type-specific effects. circRNA identification is computational: a five-method consensus and cross-referencing against independent catalogues reduce false positives, but individual events are not validated experimentally. Every paired normal comes from a tumour-bearing animal, so an effect of exposure cannot be separated from a systemic effect of carrying tumours; that would need DEN-treated animals that did not develop tumours. Finally, the design is cross-sectional: the paired normals show that the post-transcriptional state does not require transformation, but whether it contributes to transformation can only be settled by perturbation.

## Acknowledgements

We thank the Liver Cancer Evolution Consortium for generating the mouse liver cancer resource on which this study depends, and for releasing it in full. The cohort, its histopathological classification and the sequencing analysed here are described in Aitken et al. (2026) and deposited under accessions E-MTAB-16391, PRJEB15138, S-BSST383 and S-BSST384. We thank Mauro Santibanez-Koref (Newcastle University) for comments on an earlier draft of this manuscript and for his support of this work over many years.

We thank the Cancer Research UK Cambridge Institute core facilities and the EMBL-EBI Technical Services Cluster, which generated and supported the underlying data. Elements of this work were presented at EMBL-EBI and the Wellcome Sanger Institute on 3 March 2022. Initial analyses were performed at EMBL-EBI. Computational resources for the reanalysis presented here were provided by Dysplasia Diagnostics Limited, supported by a compute grant from NVIDIA Innovation Lab.

## Methods

### Cohort

The animals, husbandry, DEN treatment, tissue sampling and histopathological classification are those of the Liver Cancer Evolution Consortium cohort and are described in full by Aitken *et al.*^5,45^. No animal work was performed for this study. In brief, fifteen-day-old male mice of four backgrounds, *M. m. domesticus* (C57BL/6J and C3H/HeOuJ), *M. m. castaneus* (CAST/EiJ) and *M. caroli*, received a single intraperitoneal dose of diethylnitrosamine at 20 mg/kg, and liver lesions were dissected and diagnosed by histopathological review, with dysplastic nodules graded low or high and carcinomas graded by differentiation.

Two features of that design bear directly on what can be measured here and are stated for that reason. Sampling intervals differ by background, reflecting tumour latency: C3H/HeOuJ 22–39 weeks post-DEN (median 25), C57BL/6J 34–37 (35), CAST/EiJ 36–43 (39), *M. caroli* 56–80 (80). In C57BL/6J, histologically normal liver from treated animals was harvested at the same time as tumours from twelve animals, 34–37 weeks after DEN (median 35); two C3H/HeOuJ animals also contributed treated normal liver, insufficient for the timing analyses reported here. DEN-treated animals were male throughout; the single female in the cohort bears a spontaneous lesion, so sex is not a variable in any DEN-exposed comparison reported here. The C57BL/6J cohort additionally includes *Ctcf* heterozygous animals (HE;WT, HE;HE), and analyses requiring that class are restricted to C57BL/6J and identified as such throughout.

### Sequencing

Sequencing was performed by the Consortium and is described in full by Aitken *et al.*^5,45^. The parameters below are restated because several measures reported here depend on them. Whole-genome sequencing and ribosome-depleted total RNA sequencing were performed on matched material. Genomic DNA and total RNA were isolated simultaneously from tumours, paired normal liver and untreated normal liver using the AllPrep 96 DNA/RNA kit (80311, Qiagen); P15 liver used the AllPrep DNA/RNA mini kit^45^.

RNA concentration was measured by NanoDrop spectrophotometry and integrity assessed on a Total RNA Nano Chip Bioanalyzer (Agilent). Sequencing libraries were generated from 1 µg total RNA using the TruSeq Stranded Total RNA Library Prep Kit with Ribo-Zero Gold (20020596 and 20020599, Illumina), quantified by qPCR (Kapa Biosystems) and sequenced paired-end on an Illumina HiSeq 4000, with an average of over 26 million uniquely mapped reads per sample, approximately 75% of the reads sequenced. Most libraries were sequenced at 75 bp; a minority were sequenced at 150 bp or across both run types, and read length is recorded per sample. The animal-level intraclass correlation reported for C3H/HeOuJ, the background where both run types are common, was recomputed within the 75 bp libraries alone and is unchanged (0.38, *P* = 0.019, 117 libraries from 45 animals). That figure is STAR’s uniquely mapped count; reads mapping to up to ten loci were additionally retained in the alignments (see Alignment and expression quantification).

Although the libraries are stranded, RegTools junction extraction was run with -s 0 and rMATS with -- libType fr-unstranded. In both analyses feature identity and strand are defined from the genome and the GTF rather than from the library protocol: RegTools resolves each extracted junction through junctions annotate, and rMATS enumerates events from the GTF before any read is counted.

For SEI, annotated junction strand is subsequently enforced directly when overlapping BAM reads are selected (see *Splicing efficiency*). For rMATS, fr-unstranded means read orientation is not used as an additional counting filter: junction reads are assigned to GTF-defined events by genomic coordinates. This may admit antisense-compatible reads at loci sharing relevant coordinates, but the same counting rule is applied to every sample and background, so the contrasts reported here compare like with like. Whole genomes were prepared with TruSeq PCR-free libraries and sequenced paired-end at 150 bp on an Illumina HiSeq X Ten to a minimum of 40× coverage^5^.

### Reference genomes and annotation

Reads were mapped to the reference assembly for each background, with genic annotation from Ensembl v91^45,52^ (release 102 for circRNA identification):

| background | assembly | accession |
| --- | --- | --- |
| C57BL/6J | GRCm38 | GCF_000001635.26 |
| C3H/HeOuJ | C3H_HeJ_v1<sup>a</sup> | GCA_001632575.1 |
| CAST/EiJ | CAST_EiJ_v1 | GCA_001624445.1 |
| *M. caroli* | CAROLI_EiJ_v1.1 | GCF_900094665.2 |

^a^The C3H assembly is named for C3H/HeJ; the animals sequenced here are C3H/HeOuJ, and the C3H/HeJ assembly is the reference used for that background throughout this cohort^45^. Using a strain-specific assembly for each background, rather than mapping all four to GRCm38, is what makes the stringent alignment settings below tenable across backgrounds that differ from C57BL/6J by up to 2.5 × 10^7^ singlenucleotide positions genome-wide^45^, a divergence comparable to that between two randomly selected humans. Backsplice coordinates from all backgrounds were subsequently projected onto GRCm38 for cross-background comparison.

Annotation depth is not uniform across these four backgrounds even at a common Ensembl release. The C57BL/6J annotation was revised approximately three times annually over the period spanned by this cohort, while the three non-reference backgrounds were revised once. Measures defined relative to annotation, NSP in particular, and the known/novel splice-site classification it derives from, are therefore not directly comparable in absolute level between backgrounds, and are reported and compared within background.

### Alignment and expression quantification

#### Splice-aware alignment

Reads were aligned to the strain-specific assembly with STAR (v2.7.10a)^85^, using --alignIntronMa× 1000000, --outFilterMultimapNma× 10 and --outFilterMismatchNmax 2. Input reads were first assessed with MultiQC^86^ and trimmed with Trim Galore, removing adapter sequence and bases with quality below 15. These alignments underlie the junction-level measures, splicing efficiency and the novel splice-site classification; circRNA identification used its own alignments (see *circRNA identification*).

The multi-mapping policy is material to both junction measures and is stated explicitly: reads mapping to as many as ten loci are retained, and reads exceeding that are discarded. Junction counts at paralogous and repeat-derived loci therefore include multi-mapped support. The mismatch filter is an absolute cap of two per read, which is stringent, and is workable across the four backgrounds only because each is aligned to its own assembly rather than to GRCm38.

#### Transcript abundance and differential expression

Abundances were quantified with Kallisto (v0.43.1)^87^ using the --bias flag, against a per-background transcriptome index compiled from coding and non-coding cDNA sequences defined in Ensembl v91^45^. Transcripts per million were generated per annotated transcript and summed across alternate transcripts of the same gene for gene-level analysis; count tables include both normalised counts and a variance-stabilising transformation. Transcript classes used for the Adarb1 analysis, retained intron and nonsense-mediated decay, were taken from the Ensembl release 103 annotation and matched to the quantified transcripts on unversioned identifiers; only the class label is taken from that release, so the change of assembly it carries does not enter any coordinate used here. Differential expression was called with DESeq2^88^ (v1.48.1) at gene level, importing per-transcript estimated counts via tximport^89^, separately for each background on the protein-coding genes of the cohort count tables^45^, with sample class (untreated P15, untreated adult, paired normal or tumour) as the only term in the design. Genes were tested where at least as many libraries as the smallest class had ten or more reads. The C57BL/6J driver-specific and paired-normal contrasts come from a model with one class per lesion condition. Gene ontology over-representation was tested separately among genes higher and lower in tumours, as for alternatively spliced genes (below), against the genes that background’s analysis tested. Contrasts reported here are per background, untreated adult liver against tumour, at adjusted *P* < 0.05; the cohort paper’s cross-background intersection uses a stricter adjusted *P* ≤ 0.005 and is a different quantity. Gene expression specificity was summarised per gene using Tau^47^, on the standard definition where 0 indicates uniform expression across samples and 1 indicates expression restricted to a single sample.

### Gene set variation analysis

Per-sample pathway activity was scored with GSVA^90^, using the MSigDB v7.4 hallmark collection^91^ (h.all.v7.4.symbols.gmt, 50 sets). Input was the four-strain variance-stabilised count matrix (14,901 genes × 509 samples), which places every background and every sample class on one scale. Mouse gene symbols were matched to the human hallmark symbols by upper-casing, the usual convention for one-to-one orthologues; the median set matched 84.6% of its members, and the three weakest were PANCREAS_BETA_CELLS (65%), KRAS_SIGNALING_DN (70%) and ALLOGRAFT_REJECTION (74%). Scoring used a Gaussian kernel CDF with the mx_diff statistic and a minimum set size of 15.

Scores were recomputed for the whole cohort: median Spearman ρ per hallmark set of 0.964 (C57BL/6J), 0.974 (C3H/HeOuJ), 0.965 (CAST/EiJ) and 0.958 (*M. caroli*), with 90–94% of sets above 0.9. All GSVA results reported here come from the recomputed table.

### Variant calling, signatures and tumour composition

Variant calling and the derived mutational quantities are the Consortium’s and are described in full by Aitken *et al.*^5,45^. They are restated here because tumour cellularity and the DEN signature exposures are used as covariates and predictors throughout this manuscript. Whole genomes were aligned with bwa-mem (v0.7.12)^92^ to the strain-specific assembly, with the *S. cerevisiae* S288c reference (GCA_902192305.1) included among the alignment targets to account for spike-in DNA^45^. Regions of abnormal coverage, identified from non-tumour liver, ear and tail samples sequenced contemporaneously, were masked (5.5% of the reference in C57BL/6J, 12.7% C3H/HeOuJ, 11.5% CAST/EiJ, 12.5% *M. caroli*)^45^.

Single-base substitutions were called with Strelka2 (v2.8.4)^93^, with fixed genetic differences and segregating germline variants within the colonies identified and discarded^5^. Indels were filtered on the same parameters with two additional criteria: at least three independent supporting reads, and removal of any indel found in at least two tumours from the same animal and shared by more than half of that animal’s tumours, an animal-level filter that removes germline indels surviving the familial clustering filter^45^. Consequences were annotated with Ensembl VEP (v109.3)^94^ against the Ensembl v91 annotation for the corresponding assembly.

#### Mutational asymmetry and lesion segregation

Genomic segments with fewer than 100 mutations, or with significantly atypical read depth, were removed. Segments were classified by the absolute mutational asymmetry parameter |S| as symmetric (<0.2), asymmetric (>0.8) or intermediate. Tumours with fewer than 10% asymmetric autosomes were defined as non-asymmetric; all others asymmetric^45^.

#### Cellularity

Variant allele fraction was computed as (1 − R/d), where R is the reference read count at a mutated site and d the total depth. The modal VAF per tumour was taken as the largest peak of the VAF density (R package modes v0.7.0, function amps). Cellularity was then calculated as ploidy × modal VAF, with ploidy set to 2 for all samples except *M. caroli* tumours with no mutational asymmetry, which were inferred to be whole-genome duplicated and assigned ploidy 4^45^. This construction matters for how cellularity may be used. It is a derived quantity, not a measurement of tumour cell content, and its ploidy term is assigned by the whole-genome-duplication inference itself. In any comparison whose grouping variable is duplication status, cellularity therefore embeds the exposure and cannot serve as an independent covariate; this is why the duplication contrast reported in the Results is unadjusted. This does not apply to comparisons across groups that all carry ploidy 2, where the ploidy term is constant and cellularity varies only through modal VAF.

#### Large copy-number variants

Tumour genomes were segmented on change in read depth relative to untreated adult liver (n = 11 C57BL/6J, 11 C3H/HeOuJ, 7 CAST/EiJ, 14 *M. caroli*) with CNVkit (v0.9.6)^95^, minimum segment size 1 Mb. Copy number was derived from the log2 depth fold change FC, sample ploidy p and cellularity c as CN = ((2^FC^ − 1 + c)/c) × p, with within-strain adjustment for read-depth bias^45^. Per-sample records used here include SNV and indel counts, sister chromatid exchange counts, DEN signature exposures (den1sig, den2sig), spontaneous signature exposures, cosine signature fit, fraction of asymmetric substitutions, known driver status, fraction of segments with allelic imbalance, clone type, cellularity and VAF skew.

Single-base-substitution mutational signatures, from which the per-sample den1sig and den2sig exposures are taken, were defined and deconvolved *de novo* with the R NMF package (v0.28)^96^ using the Brunet algorithm at rank 5 over 1,500 runs; rank was selected at the first substantial drop in the cophenetic coefficient^45^. Candidate driver genes were identified with OncodriveFML (v2.2.0)^97^, OncodriveCLUSTL (v1.1.3)^98^ and dNdScv (v0.1.0)^99^ in a pan-background analysis, and a tumour was annotated as driver-positive where it carried a moderate-or high-impact mutation in an identified driver gene^45^. Clone type, recorded per lesion in the cohort descriptors as clonal, polyclonal or whole-genome duplicated, is used here as supplied and is defined with the rest of the lesion annotation in the cohort description^45^.

### Splicing efficiency (SEI)

#### Junction extraction

Splice junctions were extracted from each coordinate-sorted BAM with RegTools (v0.5.2)^100^ regtools junctions extract, run as -s 0, unstranded mode, so the library’s strand information is not used at this step. Junction strand and gene assignment come instead from regtools junctions annotate, which resolves each junction against the genome sequence and the GTF. All other parameters were at their defaults: minimum anchor length 8 bp on both sides, minimum intron 70 bp, maximum intron 500,000 bp. The 8 bp anchor requirement therefore applies to whether a junction enters the list at all; introns longer than 500 kb are outside the measure’s scope. Junctions were then annotated against the genome FASTA and the Ensembl v91 GTF for the background with regtools junctions annotate, which assigns each junction to a gene and classifies its donor and acceptor as annotated or novel.

#### Index computation

SEI was computed from the annotated junction table and the BAM by compute_sei.py, using pysam. Junctions were first filtered on the BED score column, the number of reads supporting the junction, retaining score > 1, that is, junctions with at least two supporting reads. Donor and acceptor coordinates were extracted separately and the genomic size of each junction recorded. Both the donor and the acceptor coordinate of every retained junction were queried, so each junction contributes two sites. For every site, reads overlapping the coordinate were fetched and selected by alignment orientation against the junction’s annotated strand: forward-aligned reads at a + junction, reverse-aligned reads at a − junction. Selected reads were then classified on their CIGAR string, a read containing an N operation, a skipped reference block, counted as spliced and a read without one as unspliced. This gives a spliced and an unspliced count at each donor and each acceptor, and a site-level ratio from them; where two or fewer reads overlap a site that ratio is recorded as zero, which affects the site table only, since the gene-level index is built from summed counts rather than from averaged site ratios.

Site-level counts were then grouped by gene identifier and summed, so that the gene-level index is a read-weighted ratio across all of a gene’s junctions rather than an average of per-site ratios:

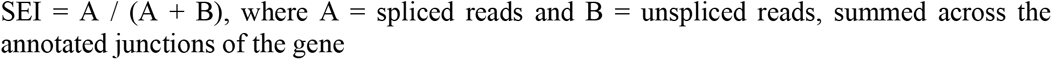

Unspliced reads at a junction originate from nascent, truncated, incompletely processed or alternatively spliced transcripts, including retained introns and exons processed at cryptic sites. Two per-sample summaries are used. The first is the mean across genes with a non-zero index, on a percentage scale and around 83–89; it is the quantity reported for splicing efficiency throughout the Results and in Figs. 2 and 7. The second is the mean across all genes, including those with no spliced read at an annotated junction, on a 0–1 scale and around 0.56; it is used in the analyses of relationships between measures and the intraclass correlations. Per-sample indices and the splice-signal scores are supplied as Supplementary Data 2.

#### Three properties of this definition bear on interpretation

First, the reads counted at a site are those overlapping the queried donor or acceptor coordinate, and a read is classified as spliced when its CIGAR string contains a skipped block, wherever in the read that block falls. A junction enters the list only with an 8 bp anchor on both sides, and no overhang is required of the reads counted at it. Second, junction strand and gene assignment come from regtools junctions annotate, which resolves each junction against the genome sequence and the GTF, rather than from the library, since extraction is unstranded (-s 0). Third, strand is enforced at two points rather than none. The strand of each site is the one regtools junctions annotate assigned from the splice motif and the GTF, so it is annotation-derived rather than inherited from the aligner or the library; read selection at the BAM then enforces that strand directly, retaining forward-aligned reads at a + site and reverse-aligned reads at a − site. A consequence is that, for a paired-end library whose mates align in opposite orientations, one mate of a properly paired fragment is counted at any given site, so the index rests on a consistent subset of the overlapping reads rather than on all of them. All three are fixed by the operational definition and applied identically to every sample and background, so the comparisons reported here are made on a common index.

#### SEI variation

For each gene, SEI variation is the standard deviation of SEI across the tumours of a background. Expression specificity (Tau^47^, in tumours) was regressed on SEI variation across genes by ordinary least squares, separately for each background, and the variance explained (R²) is reported. Both per-gene quantities are taken from the cohort’s per-gene table strain_tau_sei.tsv, which covers tumours only.

### Novel splice proportion (NSP) and alternative splicing

regtools junctions annotate classifies the donor and acceptor of each observed junction as annotated or novel against the Ensembl v91 GTF for the background, giving three classes: known, where both donor and acceptor are annotated; completely novel, where neither is; and partially novel, where one is annotated and the other is not, as in a known donor paired with an unannotated acceptor. This classification is the basis of the novel splicing measures, and it inherits the extraction thresholds above, an 8 bp anchor, and at least two supporting reads after the score > 1 filter.

NSP is the proportion of splice junctions in a gene that are cancer-specific, defined as junctions observed in tumour samples and absent both from annotation and from normal samples of the same background. Splice signal strength at donor and acceptor regions was scored with MaxEntScan^56^ for both annotated and cancer-specific sites. Alternatively spliced events were identified with rMATS turbo v4.1.1^57^, run with -- allow-clipping and --variable-read-length, -t paired for paired-end reads, --libType fr-unstranded, -- tophatAnchor 10 and --cstat 0.01. fr-unstranded is the tool’s default, so event definition remained GTF- and junction-coordinate-based while read orientation was not used as an additional counting filter; events are tested against a null of no more than 1% inclusion difference. The default statistical model was used; -- paired-stats was not set, so replicates are treated as independent rather than matched.

The retained events for every background are supplied as Supplementary Data 3. Events were taken from the junction-count (JC) output rather than the junction-plus-exon-count (JCEC) output: JC counts only reads spanning the junction, so it is not sensitive to differences in exon-body coverage, which differ substantially between tumour and normal liver here. Events were retained where the FDR was below 0.05 and the absolute inclusion-level difference exceeded 0.1, the “>10% splice-rate deviation” reported in the Results. The reported counts were generated directly from the archived JC outputs using this rule. Because NSP is defined against annotation, and annotation depth differs between backgrounds at a common Ensembl release (see *Reference genomes and annotation*), NSP is compared within background rather than across.

#### Gene-set enrichment

Genes carrying at least one retained rMATS event were tested for gene-set enrichment separately within each background, by one-sided hypergeometric test. The universe was the set of genes for which rMATS tested at least one event in that same comparison, not all annotated genes: the testable-gene universe controls for differential opportunity to detect an alternative-splicing event, which is not uniform across annotated genes. The same procedure and universe convention were applied to the reproducible tumour-only circRNA host genes of the circRNA section, where the universe is instead every gene observed to host a circRNA in any background and condition.

Hallmark, KEGG and Gene Ontology biological-process collections were downloaded once from Enrichr and the test run locally, rather than submitting gene lists to the web service, so the result does not depend on what that service was serving on a given day. The exact files are distributed with the analysis code under genesets/, with set counts and checksums in genesets/PROVENANCE.json: MSigDB_Hallmark_2020 (50 sets), KEGG_2019_Mouse (303) and GO_Biological_Process_2023 (5,406). Sets with fewer than five members inside the universe were not tested. *P* values were adjusted within each library and background by the Benjamini-Hochberg procedure, and terms reaching *q* < 0.05 are reported (Supplementary Data 1).

Gene symbols were upper-cased on both sides before intersection. The Hallmark and Gene Ontology collections are curated on human symbols while rMATS emits mouse symbols, and matching them as written recovers almost nothing: hallmark sets of two hundred genes intersect the universe at ten, and Gene Ontology returns no terms at all. Upper-casing matches orthologues sharing a symbol, which is the majority but not all of them, so genes whose mouse and human symbols differ are not counted; the median set size within the universe is reported alongside each result as the check that the intersection succeeded.

### circRNA identification (ecircscreen)

Backsplice junctions were identified with ecircscreen (v0.1.0, github.com/ddxl-osagie/ecircscreen), a pipeline that pools predictions from five published circRNA callers run over three aligners. Its design goal is to mitigate the aligner-specific biases and artefact classes that benchmarking has shown to dominate the differences between individual circRNA callers^60,64,101^.

Genome index files were built with STAR (v2.7.11b), BWA (v0.7.18) and Bowtie2 (v2.5.4)^102^, and reads aligned with each: STAR (--alignIntronMa× 1000000 --outFilterMultimapNma× 10 -- outFilterMismatchNmax 2), Bowtie2 (--very-sensitive --score-min C,-15,0 --mm -q) and BWA-MEM (-T 19). Genomes and annotation for circRNA identification were Ensembl release 102, the last release carrying GRCm38, for all four backgrounds. Predictions were pooled from five callers, each tied to a specific aligner:

| caller | aligner |
| --- | --- |
| PFv2<sup>60</sup> | STAR |
| circRNA_finder<sup>62</sup> | STAR |
| CIRCexplorer<sup>63</sup> | BWA |
| find_circ<sup>12</sup> | Bowtie2 |
| CIRI2<sup>61</sup> | BWA |

A prediction was retained only if supported by at least three of the five callers. Because no single aligner carries more than two of the five, this entails support from at least two independent alignment strategies; the corroboration-across-aligners property the filter exists to enforce is a consequence of the count threshold rather than a separate criterion. Breakpoints were required to match exactly, with no positional tolerance, and no read-support or splice-motif threshold was applied at identification, so that motif composition remained available afterwards as an independent quality measure rather than being imposed as a filter.

Retained predictions were then removed if they were observed in only one sample, overlapped segmental duplications, or had BSJ-flanking regions overlapping retrotransposons by more than 25%. Genomic overlaps were computed with intersectBed and annotateBed from BEDTools (v2.26.0)^103^. Retained predictions were cross-referenced against circBase^66^. An earlier implementation of ecircscreen was one of sixteen callers evaluated in an independent, developer-run community benchmark of circRNA detection^64^. Its filters are deliberately conservative, favouring specificity over sensitivity, the appropriate operating point for the cross-background comparison made here, in which a false-positive class shared across backgrounds would be indistinguishable from conservation.

#### Projection to a common coordinate frame

The four assemblies share neither a coordinate system nor a gene identifier namespace, so cross-strain comparison requires projection onto a single reference. GRCm38 was used throughout. C57BL/6J is native to it and required no projection, providing an internal control: any C57BL/6J junction that fails to round-trip indicates an error in the projection rather than a biological difference. Each junction retains its identifier in the assembly it was called on alongside its common-reference coordinate, so a projection failure in one background removes that background’s evidence for a junction without removing the junction.

Chain files projecting CAST_EiJ_v1 and C3H_HeJ_v1 onto GRCm38 were obtained from the UCSC GenArk assembly hubs. No chain has been published for CAROLI_EIJ_v1.1, so one was constructed from Ensembl release 102’s LASTZ net pairwise alignment of GRCm38 against CAROLI_EIJ_v1.1 (mmus_grcm38.v.mcar_caroli_eij_v1.1). A netted pairwise alignment already encodes the colinear ungapped blocks that a chain represents, making this a format conversion rather than an alignment step: each alignment block yields one chain, with the target placed on the plus strand and both coordinate frames recomputed where the alignment reports it otherwise.

Junctions were projected by their breakpoints rather than their spans. Each junction’s two termini were lifted independently as single-base features using CrossMap (v0.7.3)^104^, and a junction was retained only where both termini mapped to the same target contig in the same orientation, with coordinates then reassembled. Projecting the interval instead would require every base between the breakpoints to be alignable, and returns one fragment per aligned block rather than one junction. Contig naming was reconciled between Ensembl, GenBank and UCSC conventions by matching unique chromosome lengths rather than by assumed correspondence. Junctions whose two termini projected to different contigs were excluded from conservation analysis and retained separately as candidate inter-assembly rearrangements.

#### Cross-strain conservation

Projected consensus sets were intersected in GRCm38 coordinates, requiring exact agreement on contig, start and end. Junctions were assigned a conservation tier by the number of strains in which they were independently identified, and the conserved set was defined as those present in two or more strains.

Projection accuracy was assessed against C57BL/6J, whose coordinates are native and therefore cannot carry a projection error. Agreement between projected junctions and the C57BL/6J consensus was measured at exact coordinates and at displacements of plus or minus 1 and 2 bp; a correct projection agrees only at zero displacement, and any systematic offset would shift the maximum. Conservation tier was then tested against two catalogues independent of this pipeline, circBase mouse (mm10)^66^ and circAtlas^65^, by exact agreement on both breakpoints. Positional intersection was not used: at a median span of 8.3 kb a junction overlaps unrelated catalogued entries frequently, which inflates the apparent overlap. Support in both catalogues rises monotonically with the number of backgrounds in which a junction is detected. Splice-motif composition provided a second caller-independent measure: flanking intronic dinucleotides were read from the assembly and scored against GT/AG, GC/AG and AT/AC, with a background rate established empirically by displacing breakpoints by 500 bp and 5,000 bp within the same genomic neighbourhoods, preserving chromosome and local base composition while destroying the junction. Expression analysis was performed throughout on strain-specific identifications in each strain’s own assembly, not on the projected or conservation-filtered sets. Projection establishes cross-strain correspondence only; every quantitative measure derives from reads aligned to the assembly of the strain from which they came.

The accepted junction set, both as the projected multi-background union and as the strain-specific calls it is built from, is supplied as Supplementary Data 4. Two derived quantities are used. The abundance proportion (AP) is defined per host gene per sample as AP = BSJ / (BSJ + CSJ), where BSJ is the backsplice read count summed over the gene’s accepted junctions, quantified with CIRIquant^105^ against the accepted junction set, and CSJ is the RegTools canonical junction read count for the same gene. Both terms are read counts on the same library and require no cross-sample normalisation. AP is undefined where the canonical denominator falls below five reads. The backsplice span is the genomic distance between the backsplice donor and acceptor. Three summaries of it are used, for three different analyses, and they are not interchangeable:

- **condition-level maximal relative span**: per gene, the largest span seen in any sample of a condition group, divided by gene length. A repertoire measure: what a condition reaches, not what a typical library does (Fig. 3).
- **per-sample mean relative span**: per library, the mean across genes of that gene’s longest span divided by gene length. A sample-level central tendency, and the quantity used in the coordination and ICC analyses (Fig. 8).
- **cross-gene dispersion of maximal raw span**: per library, the coefficient of variation across genes of that gene’s longest span in base pairs. A measure of architectural heterogeneity rather than of level (Fig. 7, Supplementary Fig. 4). Per-library and per-gene span tables are supplied as Supplementary Data 5. Raw rather than length-normalised because the normalised form does not reproduce outside C57BL/6J.

#### Note on independence

AP shares its denominator with the numerator of SEI and rises mechanically when canonical splicing falls; backsplice span is a coordinate distance and carries no such dependency. Analyses of the relationships between measures use SEI, per-sample mean relative span and editing level for this reason and exclude AP and NSP. Caller versions and parameters are recorded in the run configuration distributed with the code.

Junction expression by condition. For Fig. 4, backsplice and canonical junction read counts were converted to junctions per million of total splice-junction reads in each library, summed to gene level and summarised as the median across the libraries of each sample class. The change between two classes is the log₂ ratio of those medians, computed separately for backsplice and canonical junction expression, so that genes shifting toward backsplicing are separated from genes changing in overall expression. The transcriptome-wide enrichment reported in the Results is the position of the tumour-against-adult distribution on the backsplice axis.

### RNA editing

Editing sites were identified with SPRINT (v0.1.8)^76^. SPRINT requires no SNP database: it separates editing from germline variation and sequencing error by requiring mismatches of the same type to occur in clusters, and recovers hyper-edited reads by remapping reads that fail conventional alignment. Germline variation is further limited here by aligning each background to its own assembly (see *Reference genomes and annotation*) rather than to GRCm38, so strain-specific variants are part of the reference rather than apparent mismatches. Predictions overlapping segmental duplications, homopolymer repeats and known genome variants were excluded. Hyper-edited clusters were defined as regions containing multiple editing sites separated by fewer than 10 bp, with density computed as sites per cluster length.

Per-site and per-sample editing calls are supplied as Supplementary Data 6. Three per-sample quantities are used. The editing site count is the number of sites called in that sample, and scales with library depth. The intronic fraction is the proportion of a sample’s edited sites that fall in introns; as a ratio it is more robust to library depth than the raw site count, and it is the quantity used wherever editing is compared between conditions and in its correlations with cellularity, latency, deaminase expression, hallmark programmes and immune composition. The mean editing level is the mean, over every site in the background’s editing matrix, of the fraction of reads at that site carrying the edit in that sample, with sites not called in the sample contributing zero; it is used in the analyses of relationships between measures and the intraclass correlations. Coverage and edited-read support thresholds are SPRINT defaults.

### Immune and stromal composition

Immune, stromal and other non-hepatocyte composition was estimated from RNA-seq with mMCP-counter (v1.1.0)^78^, on the variance-stabilised expression matrix and against the GRCm38 signature set.

### Statistics

Group comparisons use two-sided Mann–Whitney (Wilcoxon rank-sum) tests unless stated. Correlations are Spearman’s ρ. Where correlations are compared between groups, the difference is tested with Fisher’s z transformation, since one group reaching significance and another not does not establish that the two differ. Correlations between measures are computed within condition. Pooling conditions can produce estimates near zero where within-condition relationships have opposing signs, and this occurs in these data: two of the three inter-measure relationships reverse between untreated and exposed tissue. Joint structure across measures was assessed by decomposition of standardised, equally weighted per-modality matrices, with variance explained reported per modality per factor. Analyses and figures reported here were run in Python 3 (pandas, NumPy, SciPy, matplotlib, seaborn) and R 4.4 (tidyverse, ggpubr, patchwork, data.table); the R environment is defined by a container image distributed with the analysis code. Where a family of related tests is reported together, *P* values are corrected by the Benjamini-Hochberg procedure and both the nominal and adjusted values are given.

#### Animal-cluster bootstrap

The intraclass correlations reported above show that lesions from the same animal are not independent, materially so in C3H/HeOuJ and *M. caroli*. Principal lesion-level contrasts were therefore repeated by cluster bootstrap with 2,000 resamples in which animals, rather than lesions, were sampled with replacement: all samples from a drawn animal entered or left a resample together. The union of animals contributing to either group was resampled once per draw and split by group afterwards, rather than resampling the groups separately, because an animal may contribute to both: every C57BL/6J animal providing a carcinoma also provides dysplastic nodules, and resampling those groups independently would break the within-animal pairing. Percentile 95% confidence intervals for the resampled difference in group medians are reported. An untreated animal contributes a single library, so the procedure reduces to ordinary resampling for those groups. This was applied to the splicing-efficiency, intronic-editing, backsplice-span-dispersion, progression, NSP and clone-type contrasts (Supplementary Data 1). Secondary tests not covered by it are identified as lesion-level analyses and reported as descriptive, with the claims resting on them made within background.

#### Within-animal distances

The three per-sample summaries used in the coordination analysis were standardised across the C57BL/6J libraries, and each lesion placed at its Euclidean distance from its own animal’s paired normal. Editing enters as the intronic fraction, because the mean editing level is quantified from a different site table in each condition and is not comparable between them. The same distances under the Mahalanobis metric, on the pooled lesion covariance, rank the lesions almost identically (Spearman ρ = 0.87). Drivers were compared by standardising each lesion against the other lesions of its own animal and permuting driver labels within animals; lesions carrying more than one known driver, or none, are excluded from that comparison.

### Data and code availability

All accessions below are for the cohort as a whole; this manuscript analyses the RNA sequencing and, for the mutational quantities, the whole genomes.

| data | previously published | this cohort |
| --- | --- | --- |
| RNA-seq | E-MTAB-8518 (C3H/HeOuJ, CAST/EiJ; P15 only)<sup>5</sup> | E-MTAB-16391<sup>45</sup> |
| WGS | PRJEB37808 (C3H/HeOuJ, CAST/EiJ)<sup>5</sup> | PRJEB15138<sup>45</sup> |
| Whole-slide images | S-BSST383 (C3H/HeOuJ, CAST/EiJ) | S-BSST384 (C57BL/6J, *M. caroli*) |
| ChIP-seq | E-MTAB-11959<sup>58</sup> | E-MTAB-14454<sup>45</sup> |
| ATAC-seq | E-MTAB-11780<sup>58</sup> | E-MTAB-14144<sup>45</sup> |

Whole-slide images are additionally deposited under BioStudies accession S-BSST129. ArrayExpress accession E-MTAB-16391 is mirrored at the European Nucleotide Archive as BioProject PRJEB105369, study ERP186572; the three identifiers denote the same submission and either the ArrayExpress or the ENA form may be cited. That submission comprises 579 biological samples over 4,676 sequencing runs, all on an Illumina HiSeq 4000. ecircscreen, including the run configuration used here, is available at github.com/ddxl-osagie/ecircscreen.

The values behind every figure panel, the statistics reported in the text and Supplementary Data 1 to 6 are deposited at Zenodo (https://doi.org/10.5281/zenodo.23066938).

### Related descriptions of this cohort

The animals, treatment and sequencing are described across a series of studies of the same resource, which should be cited for full experimental detail:

- ^45^ Aitken SJ *et al.* Genetic background sets the trajectory of experimental cancer evolution. *Nature* (2026). The cohort paper, source of the assemblies, annotation release, library preparation, quantification and accessions recorded above. Cite it as the primary methodological reference for the resource.
- ^5^ Aitken SJ *et al.* Pervasive lesion segregation shapes cancer genome evolution. *Nature* 583, 265–270 (2020).
- ^44^ Connor F *et al.* Mutational landscape of a chemically-induced mouse model of liver cancer. *J Hepatol* 69, 840–850 (2018).
- Ginno PA *et al.* Single-mitosis dissection of acute and chronic DNA mutagenesis and repair. *Nat Genet* 56, 913–924 (2024).
- Nicholson MD *et al.* DNA lesion bypass and the stochastic dynamics of transcription-coupled repair. *PNAS* 121, e2403871121 (2024).
- Anderson CJ *et al.* Strand-resolved mutagenicity of DNA damage and repair. *Nature* 630, 744–751 (2024).
- Daunesse M *et al.* (2025), nuclear–mitochondrial crosstalk. Preprint.
- Connelly J *et al.* (2026), explainable machine learning relating histological to genomic pathology. Preprint.

## Supplementary figures

**Supplementary Figure 1.**
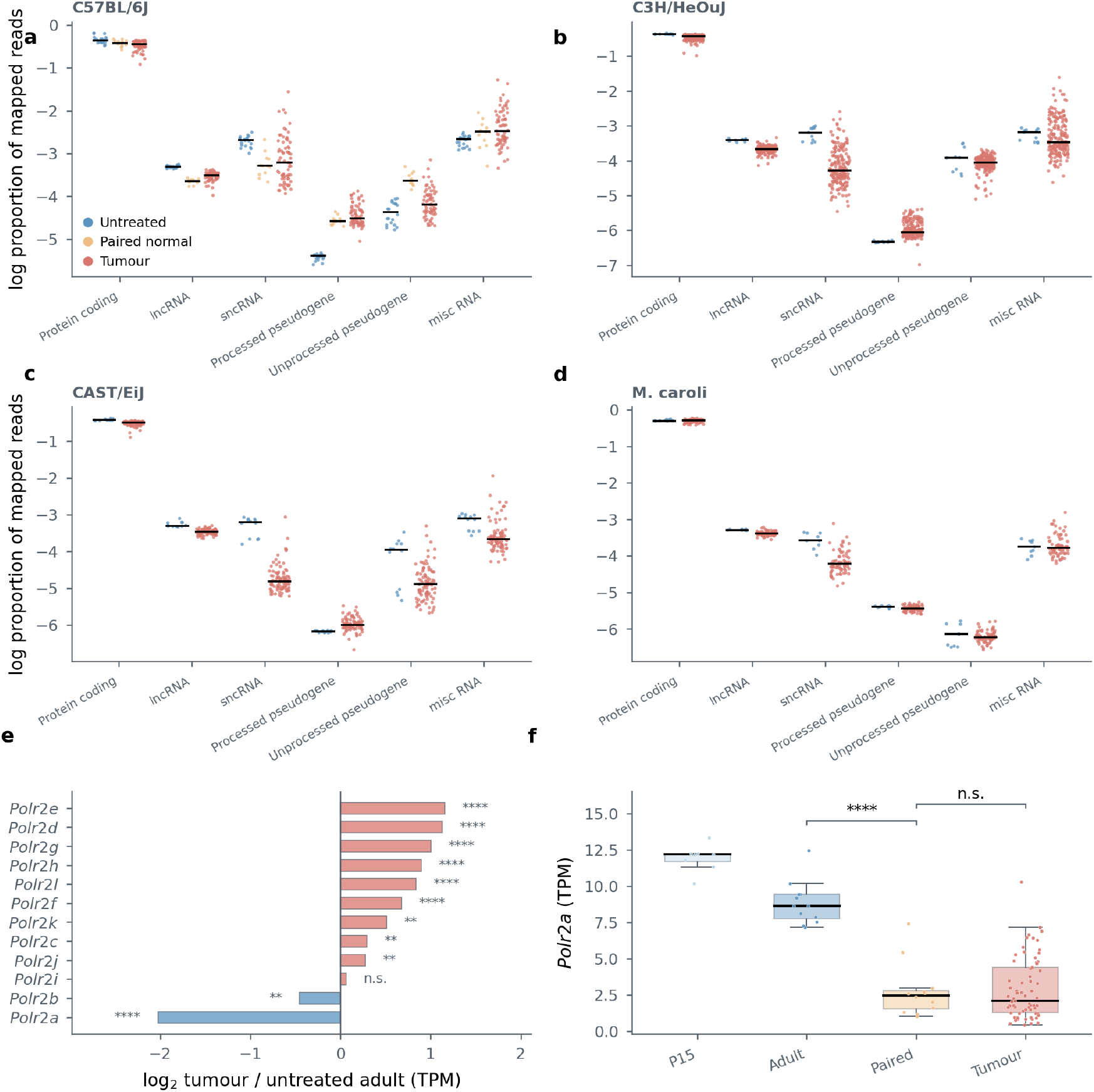
Transcriptome composition in exposed tissue. **a**-**d**, Read composition across RNA classes, one panel per background: C57BL/6J (**a**), C3H/HeOuJ (**b**), CAST/EiJ (**c**) and *M. caroli* (**d**). Each point is a library, plotted as the natural log of the proportion of mapped reads assigned to that class, with the group median marked. Protein-coding output is tightly constrained in every background while the non-coding classes disperse in tumours, most strongly for small non-coding and miscellaneous RNA, and reads from processed pseudogenes rise. Where paired normal liver exists in usable numbers, in C57BL/6J, it sits between untreated liver and tumour for several classes. **e**, Log_2_ fold change in TPM, tumour over untreated adult liver, for the twelve Pol II core subunits *Polr2a* to *Polr2l*, coloured by direction. *Polr2m* is excluded because it encodes GDOWN1, which associates with Pol II rather than constituting it. Asterisks mark two-sided Mann–Whitney tests against untreated adult liver, Benjamini-Hochberg adjusted across the twelve subunits (** *q* < 0.01, **** *q* < 0.0001); n.s., not significant. *Polr2a*, encoding the largest subunit RPB1, falls ∼4-fold while ten of the remaining eleven rise, so the complex is not scaled up or down as a unit. Five of the twelve, *Polr2e*, *Polr2f*, *Polr2h*, *Polr2k* and *Polr2l*, are the RPABC subunits shared with RNA polymerases I and III^106,107^, and they are among the larger increases, so part of the rise may reflect general polymerase demand rather than Pol II alone. **f**, *Polr2a* TPM by condition. The fall is already complete in paired normal liver and tumours sit no lower than their paired normals, placing the change in exposed tissue rather than at transformation. Boxes are quartiles with the median, individual libraries overlaid; brackets are two-sided Mann–Whitney, unadjusted across the two comparisons shown.

**Supplementary Figure 2.**
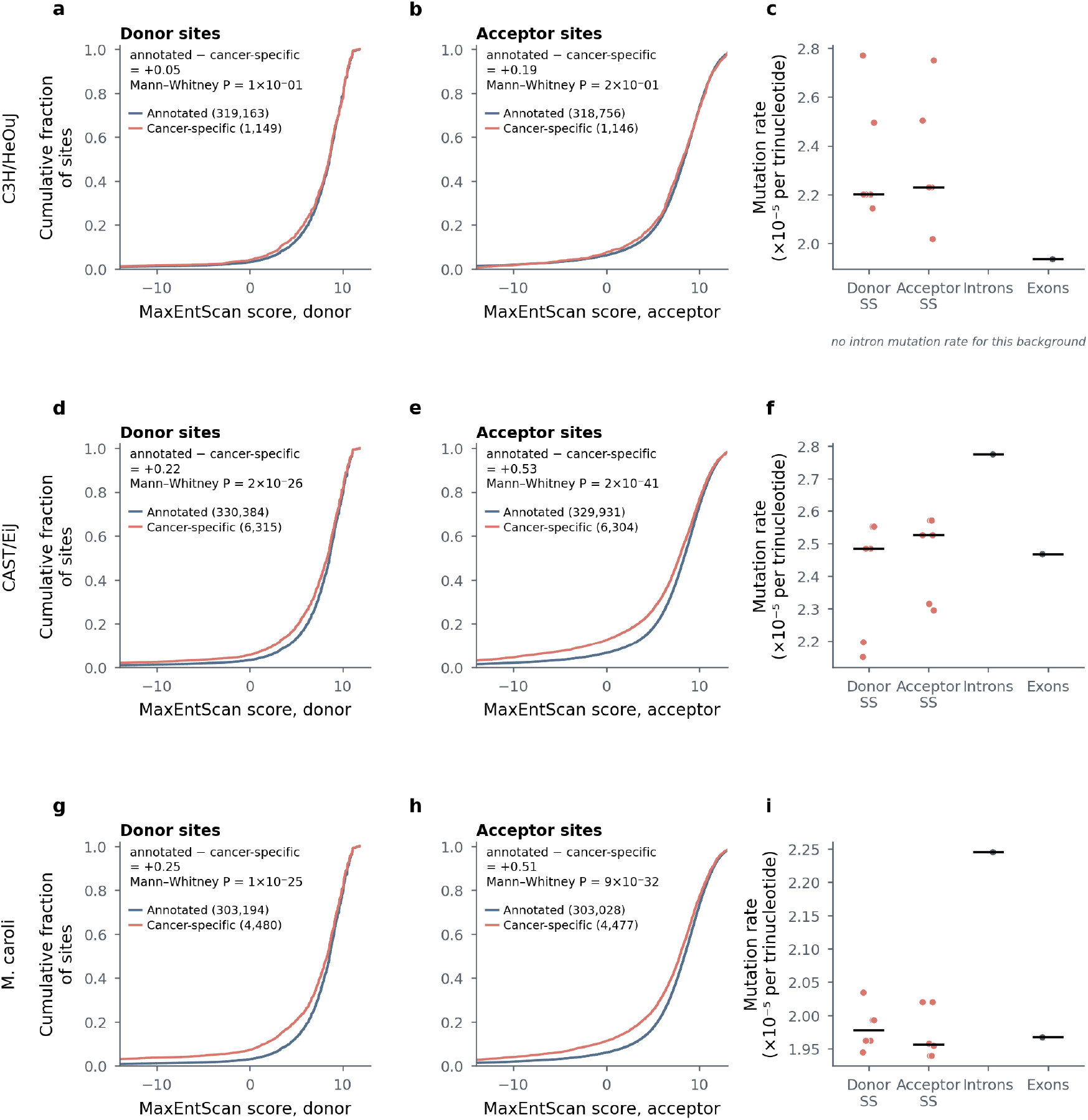
Splice-signal strength and mutation rate in the three non-reference backgrounds. Figure 2c and 2d repeated for C3H/HeOuJ (**a**–**c**), CAST/EiJ (**d**–**f**) and *M. caroli* (**g**–**i**) on the same axes. **a**,**b**,**d**,**e**,**g**,**h**, cumulative MaxEntScan distributions for annotated and cancer-specific donor and acceptor sites, with the median difference and Mann–Whitney *P*. Cancer-specific sites score lower at both site types in all three, as in C57BL/6J; the difference is significant in CAST/EiJ and *M. caroli* and not in C3H/HeOuJ, where 1,149 cancer-specific donor sites were recovered against 6,315 and 4,480. **c**,**f**,**i**, trinucleotide-normalised mutation rate by feature class; points are pooled estimates and the bar is the median. Splice sites sit at or below the intronic rate wherever the comparison can be made. No intron mutation table was produced for C3H/HeOuJ, so that class is absent there and the panel states this rather than leaving an empty column to be read as a rate of zero.

**Supplementary Figure 3.**
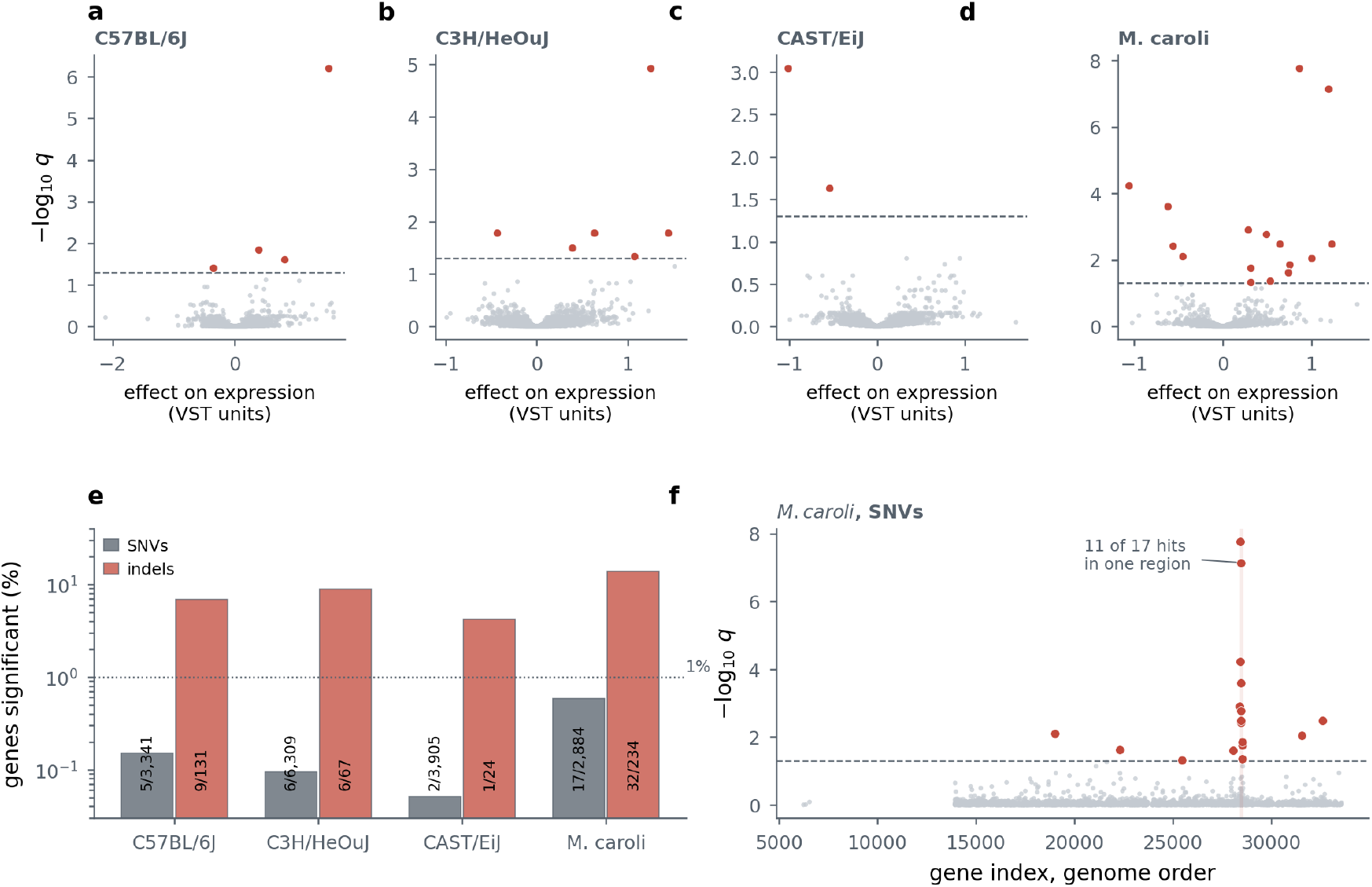
Splice-region mutations explain almost no expression variation. **a-d**, Per-gene association between carrying a splice-region single-nucleotide variant and that gene’s expression, one panel per background. Expression is variance-stabilised and each gene is fitted by ordinary least squares as expression ∼ variant present + tumour cell content + indicators for recurrent driver genes; *q* is Benjamini-Hochberg adjusted across the genes tested in that background. Dashed line, *q* = 0.05; genes reaching it are coloured. **e**, The same counts as a percentage of genes tested, for single-nucleotide variants and for indels, with the count over its denominator printed on each bar and 1% marked. Splice-region indels are rare enough that only 24 to 234 genes are testable in any background, so a handful of hits is a large fraction of a small denominator. **f**, *M. caroli* plotted against position in its annotation, which runs in genome order. Eight of the seventeen significant genes fall in a single stretch of 57 tested genes (hypergeometric *P* = 3 × 10^−10^) and eleven fall in two adjacent stretches, so this background’s apparent excess is better read as one divergent locus than as seventeen independent associations. A gene is tested where at least five libraries carry a variant and at least three libraries fall on each side of the contrast.

**Supplementary Figure 4.**
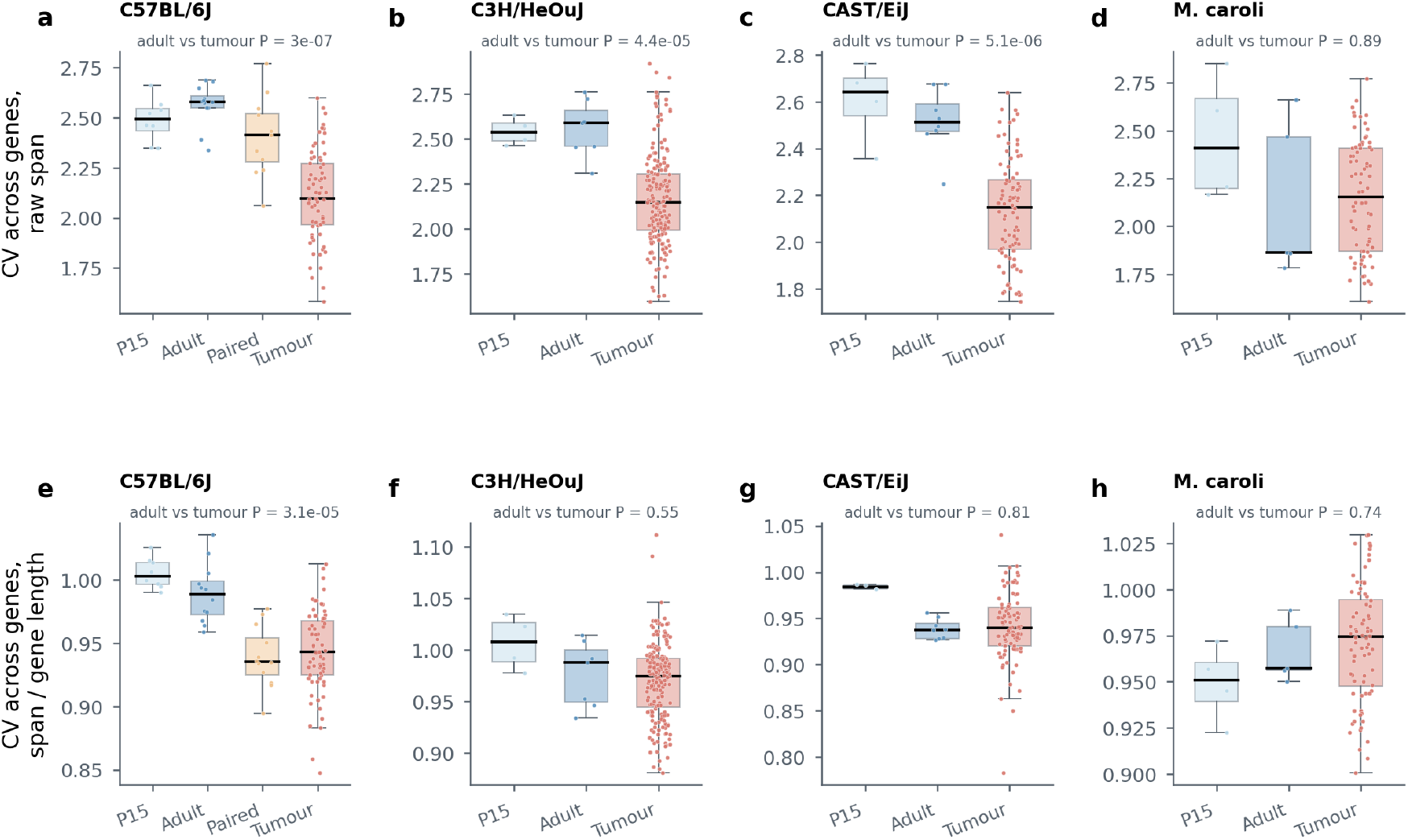
Cross-gene dispersion of backsplice span in all four backgrounds, on raw and on length-normalised distances. **a**-**d**, the statistic shown in Figure 7b: the coefficient of variation across genes of each gene’s longest backsplice junction in base pairs, by condition. The untreated-to-tumour fall reproduces in C3H/HeOuJ and CAST/EiJ at effect sizes close to C57BL/6J’s (Cliff’s δ −0.81, −0.85 and −0.93) and is absent in *M. caroli*, whose untreated baseline differs markedly from the three *M. musculus* backgrounds. **e**-**h**, the same dispersion computed after dividing each junction by its host gene’s length. In C57BL/6J this version places the paired normals at the tumour value rather than a third of the way to it, and it reproduces in no other background. Paired normal liver exists in usable numbers only in C57BL/6J, so the paired column is absent elsewhere by design. Mann–Whitney, untreated adult against tumour.

**Supplementary Figure 5.**
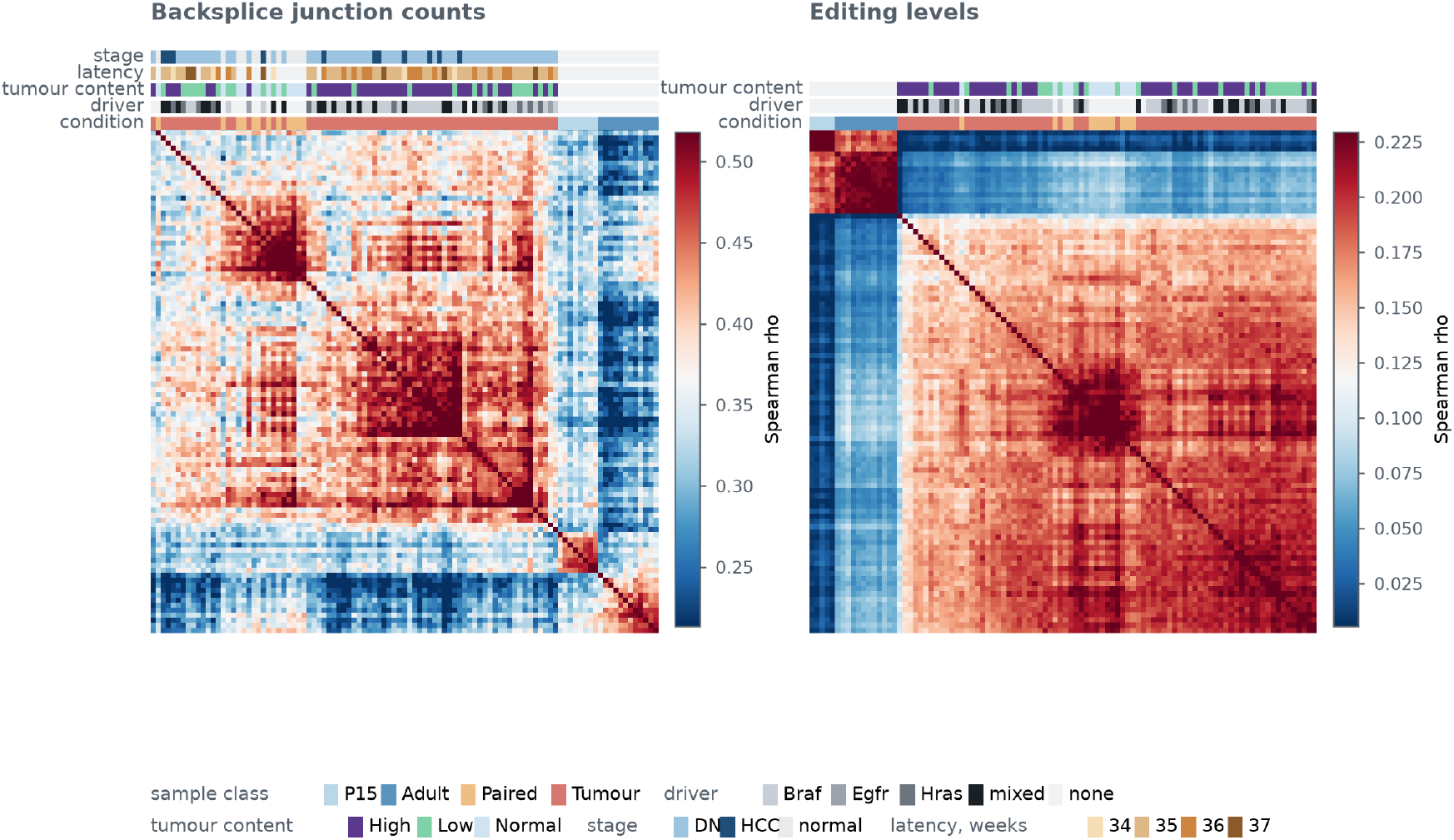
Library clustering on backsplice junction counts and on editing levels (C57BL/6J). **a**, Spearman correlation between libraries on backsplice junction counts, retained where the total across libraries exceeds the number of libraries, hierarchically clustered on 1 minus the correlation. **b**, the same on per-site A-to-I editing levels. Both on log_2_(x + 0.5). The strip above each panel mark, from the bottom, sample class, driver gene and tumour content, with latency and stage added in **a**, following the annotations of the analyses these reproduce; tumour content is thresholded at 0.5 and latency is the discrete week of sacrifice. Untreated liver separates from tumour in both measures and the two ages separate within untreated liver, while paired normal liver groups with tumour rather than with untreated liver. Colour spans the 2nd to 98th percentile of the off-diagonal correlations rather than the full range, which these occupy only a narrow part of.

**Supplementary Figure 6.**
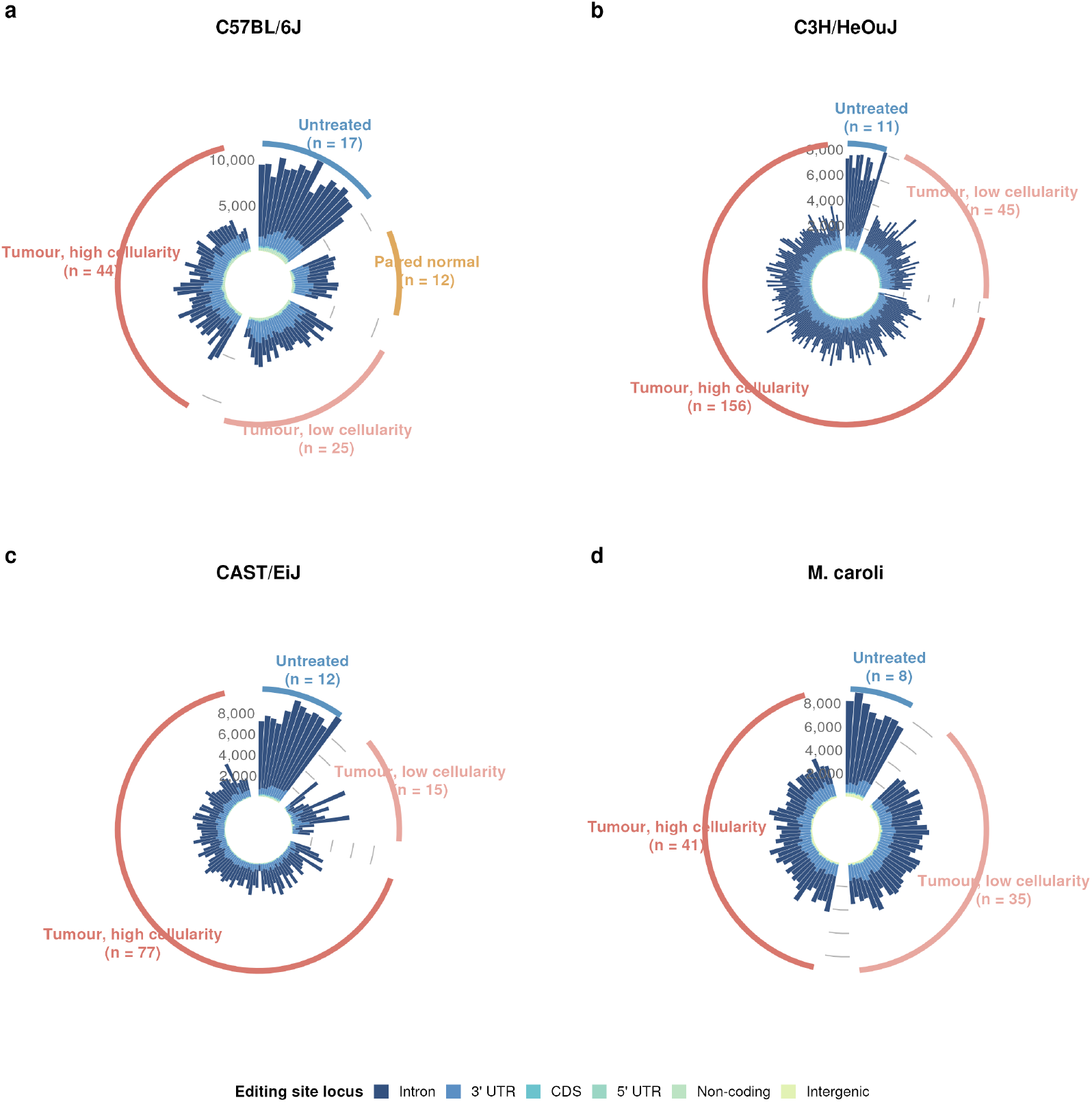
Editing site composition per sample, all four backgrounds. Circular stacked bars, one per library, coloured by the region the editing site falls in and ordered within group by intronic share, so the gradient reads around the circle. **a**, C57BL/6J. **b**, C3H/HeOuJ. **c**, CAST/EiJ. **d**, *M. caroli*. Bar height is the total number of sites called in that library and the radial gridlines give the scale. Tumours are split at 0.5 tumour cell content into high- and low-cellularity groups, which are indistinguishable, so the loss of intronic editing is not a function of tumour cell content. Only C57BL/6J has paired normal liver in usable numbers, so the paired normal group appears in **a** alone.

**Supplementary Figure 7.**
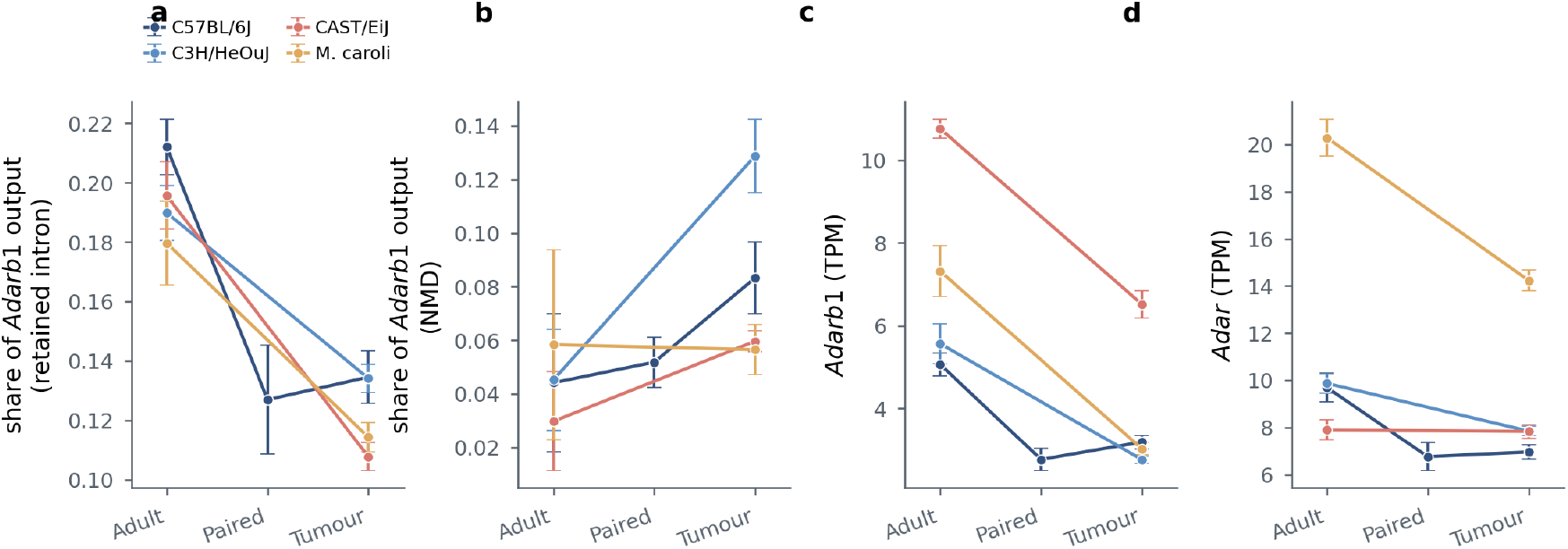
*Adarb1* transcript class composition and deaminase expression. **a**,**b**, Share of *Adarb1* transcript output carried by retained-intron (**a**) and nonsense-mediated decay (**b**) transcripts, by condition and background. Shares rather than levels, so a change means output is distributed differently rather than that the gene is transcribed more or less. **c**,**d**, *Adarb1* (**c**) and *Adar* (**d**) expression in TPM. Points are means with standard errors; the statistics quoted in the text are medians with Cliff’s δ and Mann–Whitney, which agree with these in direction throughout. Paired normal liver exists in usable numbers only in C57BL/6J, so the other three backgrounds connect untreated liver to tumour directly. Transcript biotypes are taken from each background’s own annotation cross-reference; all four annotate the same thirteen *Adarb1* transcripts in the same three classes. A shift between the two protein-coding isoforms is not shown: transcript-level quantification reports one, but neither feature distinguishing those isoforms changes, the internal 30 nt alternative acceptor sitting at 0.93, 0.91 and 0.92 across the three conditions and cassette inclusion at zero throughout, so the apparent switch is an apportionment artefact.

**Supplementary Figure 8.**
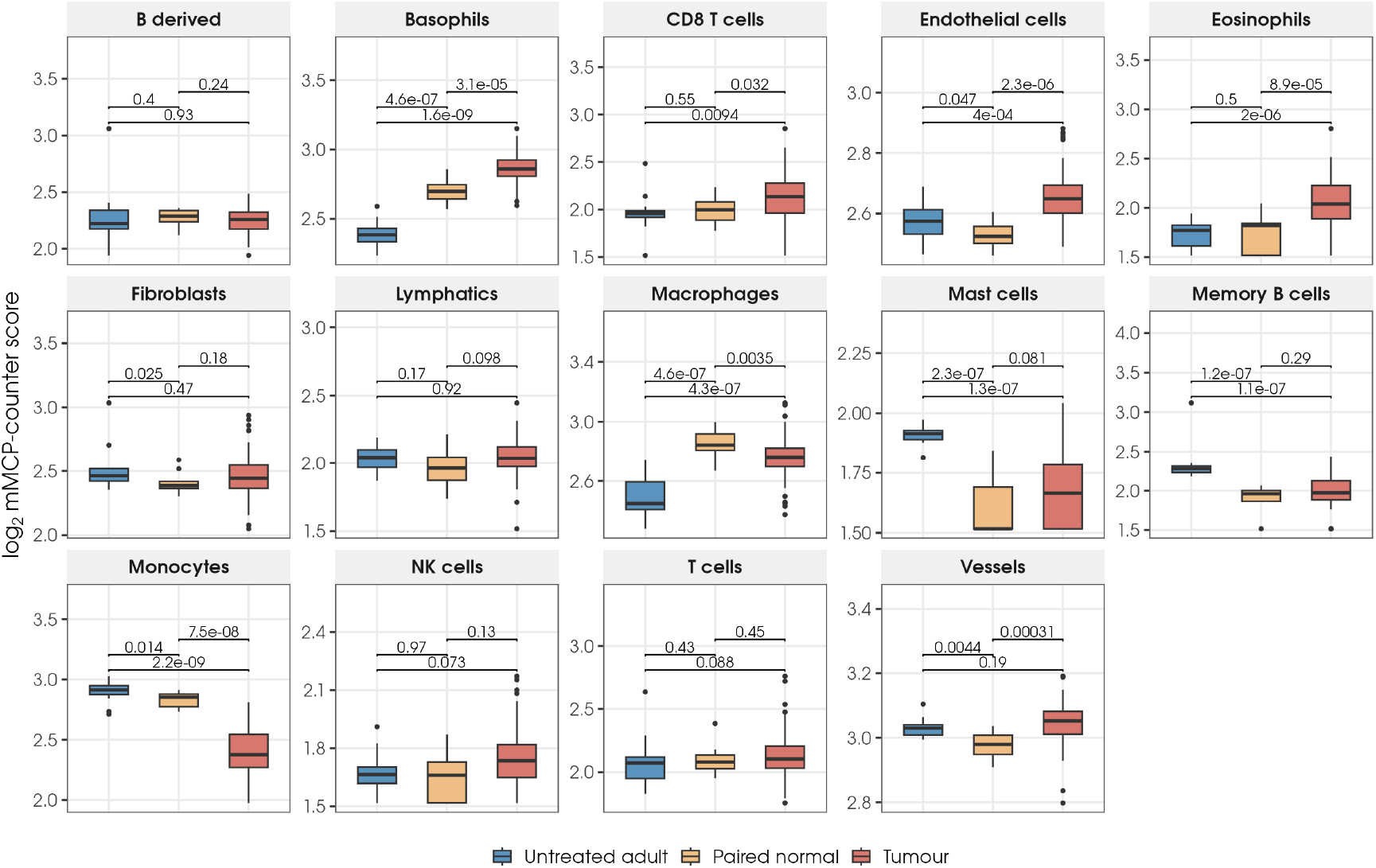
Immune and stromal composition by mMCP-counter (C57BL/6J). Scores on the variance-stabilised expression matrix for 96 libraries, by population, comparing untreated adult liver, paired normal liver and tumours. Boxes are quartiles with the median. Basophils and eosinophils are higher in tumours and monocytes lower, while macrophages, mast cells and memory B cells separate both exposed groups from untreated liver rather than separating tumours from paired normals. Pairwise Mann–Whitney, unadjusted. Neutrophils and granulocytes are absent because the signature resolves no genes for them in this genome build. Untreated P15 liver is excluded.

